# A combined test for feature selection on sparse metaproteomics data - an alternative to missing value imputation

**DOI:** 10.1101/2021.06.22.449387

**Authors:** Sandra Plancade, Magali Berland, Melisande Blein-Nicolas, Olivier Langella, Ariane Bassignani, Catherine Juste

**Affiliations:** Université fédérale de Toulouse, INRAE, UR875 MIAT, F-31326 Castanet-Tolosan, France; Université Paris-Saclay, INRAE, MGP, 78350, Jouy-en-Josas, France; Université Paris-Saclay, INRAE, CNRS, AgroParisTech, PAPPSO, doi.org/10.15454/1.5572393176364355E12, F-91190, Gif-sur-Yvette, France; Université Paris-Saclay, INRAE, CNRS, AgroParisTech, GQE-Le Moulon, F-91190, Gif-sur-Yvette, France; Université Paris-Saclay, INRAE, AgroParisTech, Micalis Institute, 78350, Jouy-en-Josas, France

## Abstract

One of the difficulties encountered in the statistical analysis of metaproteomics data is the high proportion of missing values, which are usually treated by imputation. Nevertheless, imputation methods are based on restrictive assumptions regarding missingness mechanisms, namely “at random” or “not at random”. To circumvent these limitations in the context of feature selection in a multi-class comparison, we propose a univariate selection method that combines a test of association between missingness and classes, and a test for difference of observed intensities between classes. This approach implicitly handles both missingness mechanisms. We performed a quantitative and qualitative comparison of our procedure with imputation-based feature selection methods on two experimental data sets, as well as simulated data with various scenarios regarding the missingness mechanisms and the nature of the difference of expression (differential intensity or differential missingness). Whereas we observed similar performances in terms of prediction on the experimental data set, the feature ranking and selection from various imputation-based methods were strongly divergent. We showed that the combined test reaches a compromise by correlating reasonably with other methods, and remains efficient in all simulated scenarios unlike imputation-based feature selection methods.

## INTRODUCTION

Metaproteomics refers to the study of all proteins present in an ecosystem (soil, water, gut…) at a given time. It allows for the qualitative and quantitative profiling of the tremendous diversity of proteins in complex biological samples. It is the method of choice to learn about which microorganisms are doing what in a microbial ecosystem. Therefore, metaproteomics moves beyond the genetic potential addressed by metagenomics, and it is generating rising interest and new international initiatives (www.metaproteomics.org). Yet metaproteomics has long lagged behind metagenomics due to the lack of appropriate tools, but impressive progress in LC-MS/MS technologies (Liquid Chromatography coupled with tandem Mass Spectrometry) makes it possible to decipher metaproteomes in a deep, broad and high throughput manner. However, processing of metaproteomics data is much less developed than for metagenomics and statistical approaches developed for proteomics of single organisms cannot necessarily be transposed to complex ecosystems. Indeed, metaproteomics data are characterized by a huge diversity and specificity within and between samples; this generates large and sparse matrices of protein abundances which require dedicated analytical methods. In particular, selecting metaproteomic features that are shared by homogeneous clinical groups could facilitate the diagnosis or prognosis of a disease.

Feature selection methods (FSMs) can be classified in two categories. Wrapper and embedded methods make use of a classifier to select a set of features based on their discrimination ability, either with a recursive selection (wrapper) or by including a filtering into the classifier (embedded) (Saeys et al., 2007). While these methods enable the extraction of a reduced list of predictors, they are pointed out as potentially generating overfit (Saeys et al., 2007), and lead to the elimination of correlated features which may be detrimental to biological intepretation. In univariate methods, features are examined separately. These methods do not account for potential interactions amongst variables, but they enable the inclusion of more complex designs (batch effects, multiple effects, censoring,…).

One of the difficulties encountered to implement FSMs on shotgun proteomics and metaproteomics is to handle the missing data. Indeed, LC-MS/MS technologies are known to generate a high rate of missing values and this phenomenon is enhanced in metaproteomics. On one hand, microbiota composition is largely specific to individuals, leading to a significant proportion of truly missing proteins. On the other hand, the high complexity of microbiota samples makes data acquisition and pre-processing particularly sensitive, and generates a higher technical variability than observed on proteomics data, leading to important measurement errors as well as missing values. The processes leading to missingness are diverse and may originate from any step of the pipeline, either biochemical, analytical or bioinformatics (Lazar et al., 2016). These mechanisms can be analysed in the framework developped by Rubin (1976), who distinguishes Missing At Random (MAR) in which the probability for a feature to be missing is independent of its true abundance, and Missing Not At Random (MNAR) in which missingness depends on the abundance, including notably thresholding due to device detection limit. It is commonly recognized that both MAR and MNAR occur with LC-MS/MS technologies (O’Brien et al., 2018; Lazar et al., 2016), but neither the proportion of each mechanism on a data set nor the precise mechanism at the origin of a given missing value are known a priori.

Methods to address missing data in proteomics mostly rely on either missing value imputation (Wang et al., 2020) or statistical modelling of censoring mechanisms (Karpievitch et al., 2009; Luo et al., 2009; O’Brien et al., 2018), even if a few alternative have arisen. Borrowing from both above mentioned categories, Berg et al. (2019) have recently proposed a multiple imputation approach based on a MAR/MNAR model. Gianetto et al. (2020) (R package `imp4p`) developed a statistical model that combines MAR and MNAR missing value imputation. Besides, Webb-Robertson et al. (2010) developed a filtering approach that circumvents missing values imputation by means of two successive filtering based on difference in terms of peptide occurrence, and difference of intensities among the non-missing observations. But to our best knowledge, in the metaproteomics context, the treatment of missing values mostly relies on imputation (Tang et al., 2020b). A large number of imputation methods for proteomics or metaproteomics have been proposed in literature (R package `NAguideR`, Jin et al. (2021)), and can be classified in three categories (i) single value imputation, where missing intensities are replaced by the same value for all samples; (ii) global structure methods, in which imputation is based on correlations between the whole set of observations; (iii) local similarity imputation, based only on the most similar features.

In this paper, we propose an approach which circumvents the limitations of missing value imputation and implicitly handles both MAR and MNAR mechanisms. This univariate feature selection method combines a presence/absence test which detects if the frequence of missingness is different between classes, and a test of the difference in observed intensities between classes, embedded in a permutation test procedure. We compared our method with three imputation-based FSMs, on two metaproteomics data sets: the first one from human gut microbiota of a cohort of coronary artery disease patients, and the second one from gut microbiota samples of pigs repeatedly measured in a diet perturbation experiment. Moreover, we made use of a set of technical replicates to explore missingness mechanisms.

## MATERIAL AND METHODS

### Experimental data sets

#### ProteoCardis

We used a subset of the data set generated in the ProteoCardis project, an association study between the human intestinal metaproteome and cardiovascular diseases (Bassignani, 2019, Section 1.6). Two classes were considered: patients with acute cardiovascular disease (n=49) and healthy controls (n=50). For each of these 99 subjects, the extracted gut microbiota was fractionated into its cytosolic and envelope compartments, which were analysed separately for their metaproteome, giving a total of 198 metaproteomes. Details on metaproteomics analyses can be found in Bassignani (2019) (Section 4.1.2). The cytosolic and envelope data sets are denoted by *ProteoCardis-cyto* and *ProteoCardis-env*. In order to investigate technical variability, eight biological samples from the ProteoCardis cohort were replicated seven times each, for both their cytosolic and envelope compartments analysis.

The peptides and the proteins they come from were identified using an original iterative method described in Bassignani et al. (2021). Indistinguishable proteins, i.e. those identified with a same set of peptides, were grouped into metaproteins (or protein subroups) using the parsimonious grouping algorithm X!TandemPipeline (Langella et al., 2017). To simplify the writing, those protein assemblages are denoted ‘‘proteins” in the following (Bassignani et al., 2021; Bassignani, 2019). Finally, intensities of proteins were calculated as the sum of the extracted ion currents of their specific peptides, using MassChroQ (Valot et al., 2011). Data are available at https://doi.org/10.15454/ZSREJA.

#### Pigs

The data set *Pigs* consists in fecal microbiota analyses on 12 pigs observed at six times points during a four-week diet. Samples from weeks one and two (one observation per week) were gathered into a “metabolic period” and samples from weeks three and four (two observations per week) into an “equilibrium period”. These two periods represent our classes of interest for this analysis, similarly to Tang et al. (2020b). All details can be found in Tilocca et al. (2017), and the data are available at ProteomeXchange PXD006224.

#### Filtering of sparse features

FSMs were applied on log-transformed data after filtering out proteins with less than *τ* non-missing values, with *τ* equal to 40% of the size of the smallest class (*τ* = 20 for *ProteoCardis* and 10 for *Pigs*). This value represents a compromise between a high threshold that may lead to the deletion of a large part of the features, and a low threshold where too little information would be available for some variables. Nevertheless, as the impact of the missing value treatment may depend on the proportion of missing values, complementary analyses were performed with higher threshold values (30, 40 and 50 for *ProteoCardis* data sets; 20 and 30 for *Pigs*).

#### Statistical characteristics of the experimental data sets

Even after filtering of sparse features, *ProteoCardis* data sets are still highly sparse, with most proteins having more than half missing values while *Pigs* displays a larger proportion of proteins with very few missing values (Figure S1, top). These differences of sparsity may originate from a higher similarity in terms of genetic background and diet among experimental animals. Moreover, many more proteins are significantly different between the two classes in *Pigs* than in *ProteoCardis* for all FSMs (Figure S1, bottom). Thus *Proteocardis* and *Pigs* display different statistical characteristics, which enhance the robustness of the FSM comparison carried out in this paper.

### Analysis of replicates

Consider a technical replication of the analysis of a biological sample *i*. The probability that a feature *j* is missing in the technical replicate, given that the observed intensity *Y_i,j_* is equal to y in the original analysis, is defined as follows:

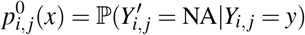

with 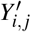 the observed intensity in the replicate. The replicate data sets were used to infer the missingness probability function under the assumption that the probability 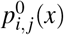 is independent of the biological sample and of the feature: 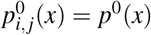. Then, observed intensities were stratified in 5% quantiles: (*y*_0_,…, *y*_20_) and *p*^0^ was approximated by:

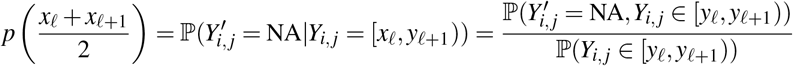

which was estimated by its empirical counterpart:

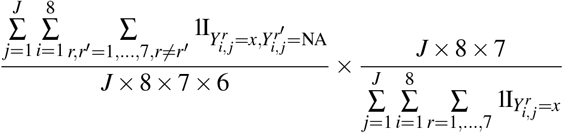

with 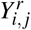 the intensity of the protein *j* in the replicate *r* of the biological sample *i*.

### Simulation study

A simulation study was conducted to illustrate the impact of both the nature of the biological difference between classes and the missingness mechanism. A realistic full data set was generated by kNN imputation of missing values on the log-transformed intensities from the *ProteoCardis-cyto* data set, after filtering of proteins with less than 10 non-missing values. Two classes of size 49 and 50 were randomly sampled among the 99 samples, so that no proteins are differentially expressed between the two classes except by chance. Then, difference of expression between classes was generated. First of all, 2000 proteins were randomly selected to be truly differentially expressed, assuming either a difference in intensity (fold change), or a difference in probability of presence. Then, missing values were picked up assuming either MAR or MNAR mechanism, followed by filtering of proteins with less than 20 non-missing values. Details can be found in Supplementary Material.

### Combined test

We propose a protein level univariate combined test that accounts for both missing and non-missing data. Consider *m* features (here, proteins) observed in *n* samples belonging to two or more classes. For each feature *j*, the difference of intensity between the two classes on non-missing observations, and the association between class and missingness are tested via the following linear mixed model (lmm) and generalised linear mixed model (glmm):

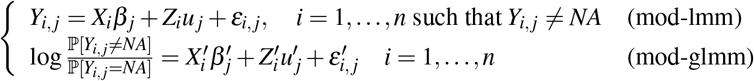

with *Y_i,j_* the log-transformed observed intensity of feature *j* in sample *i*, (*X*, *X′*) design matrices of fixed effect including the class effect, (*Z*, *Z′*) design matrices of random effects, (*ε_i,j_*)_*i*_ and 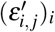 i.i.d. centered gaussian. Let *p*_1_ and *p*_2_ be the *p*-values of the F-test for class effect in models (mod-lmm) and (mod-glmm) respectively.

Let *S* be the Fisher combined statistic defined as:

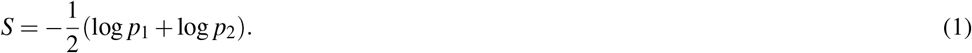

*S* is large if at least one of the two *p*-values *p*_1_ and *p*_2_ is small; Moreover, if only one of the two *p*-values is small, *S* is weakly affected by the value of the largest one. If the statistics of the two tests were independent, *S* would be *χ*^2^-distributed under the global null hypothesis, but this assumption may be violated, especially under MNAR assumption, since low protein abundances could simultaneously lead to low observed intensities and high probability to be missing. Therefore, the distribution under the global null hypothesis is obtained by repeated permutations (*N^perm^*) of the classes. In order to reduce the computing time and to increase the number of distinct values that can be taken by *S*, the distribution under the null hypothesis is assumed to be identical for all variables with the same proportion of missing values. Mathematical details are provided in Supplementary Material.

#### Design for ProteoCardis and Pigs data sets

For the *ProteoCardis* data sets, no random effect was considered and

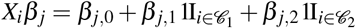

with 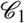 and 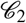 the two classes. For the *Pigs* data set, a random animal effect was added:

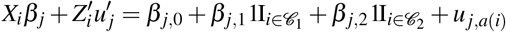

with *a*(*i*) the animal on which sample *i* was collected. Comparison between classes is performed based on the contrast *β*_*j*,1_ – *β*_*j*,2_.

#### Permutations framework

In the case of a complex design, single shuffling may be inappropriate since data are not freely exchangeable under the null hypothesis. Thus for *Pigs*, permutations were implemented such that the number of observations on each animal over each period was preserved (two time points and four time points over the first and second period respectively), using the R package `permute`.

For the implementation on the whole data set, we considered *N^perm^* = 10^5^, and for the prediction accuracy and the replicability on independent subsets, *N^perm^* = 10^4^. A larger number of permutations leads to a better precision of the *p*-values (of order 1/*N^perm^*) but at the cost of a larger computation time which increases linearly with *N^perm^*. Note that the procedure can be parallelised very easily, since the distribution under the null hypothesis is computed separately for each possible number of missing values. Besides, the combined test requires a sufficient sample size, so that enough permutations can be realised to compute the statistic distribution under the null hypothesis with a good precision.

### FSMs based on NA imputation methods

The combined test was compared with feature selection procedures based on missing value imputation. We considered the three imputation methods proposed by Tang et al. (2020a) (package `metaFS`), namely (i) single value imputation, where all missing value are replaced by the smallest intensity observed in the data set; (ii) k-Nearest Neighbours (kNN); (iii) Singular Value Decomposition (SVD). Following Wang et al. (2020) kNN was implemented using the R function `SeqKNN` with a number of neighbours *k* = 10 and SVD was implemented using the function `pca` (package `pcaMethods`) with two components. Then, the linear mixed model (mod-lmm) was applied on the vectors of observed and imputed intensities. The choice of a lmm testing procedure after imputation guarantees that the differences observed between the imputation-based FSMs and the combined test are exclusively due to the treatment of missing values. The imputation-based FSMs are denoted as follows.

- **Single-lmm:** log-transformation + single value imputation + lmm
- **KNN-lmm:** log-transformation +kNN imputation + lmm
- **SVD-lmm:** log-transformation + SVD imputation + lmm

### Hurdle model

Goeminne et al. (2020) proposed a peptide level model, the Hurdle model, that presents similarities with our approach, by combining MSqRob, a mixed model applied on all peptides observed intensities of each protein including a random sample effect, and a quasi-binomial model on the number of observed unique peptides. The *p*-values are combined as in (1), but independence is assumed between both statistics. Functions to implement the tests are available at https://github.com/statOmics/MSqRobHurdlePaper.

Following Goeminne et al. (2020), additional filterings at the peptide level were applied to *Proteo-Cardis* and *Pigs* datasets prior to the Hurdle test implementation. Peptides with only one identification were deleted, as the model would be perfectly confounded. Then, proteins identified by only one peptide were removed.

### Resampling-based procedure for control of False Discovery Rate (FDR)

To account for correlation between variables, the False Discovery Rate (FDR) was controled using the resampling-based procedure proposed by Reiner et al. (2003): *p*-values were reestimated by resampling (100 times) from the marginal distribution prior to *p*-value adjustment (Benjamini and Hochberg, 1995a). Even if most FDR procedures only guarantee an upper-bound control and are subject to assumptions on dependence between variables, the number of selected variables for a given FDR threshold is an indicator of the FSM’s power. Moreover, the resampling-based FDR procedure considered here enables the bias due to complex dependence structures to be circumvented. Mathematical details are provided in Supplementary Material.

### Criteria for method comparison

#### Agreement between FSMs

The Pearson correlation between vectors of log-transformed *p*-values quantifies the overall similarity between FSMs on all proteins, while putting more weights on those with low *p*-values via the log-transformation. The proportion of common selected features among the top *N* directly targets agreement in terms of feature selection. Values of *N* were chosen for each data set according to the number of significant features. For *ProteoCardis* data sets which display a small number of significant values, we considered *N* = 30, 100, 200. For *Pigs*, as a large number of proteins are significantly different between the two classes, we considered larger values *N* = 200, 500, 1000. Moreover, for *Pigs*, sample splitting in cross-validation loop and stability computations was implemented such that all observations from the same animal remained in the same subset.

#### Stability of feature selection between independent data sets

The set of samples was repeatedly (100 times) split into two independent subsets while preserving the proportion of each class. FSMs were applied on each subset, and the stability was quantified as the proportion of common variables among the top *N* features selected on each of the two subsets.

For comparison between the combined test and the Hurdle test, we considered alternative criteria that account for the difference in the total number of features, since the number of tested proteins differs due to the additional filtering for the Hurdle model. Cohen’s kappa (McHugh, 2012) enables to compare the agreement between two “raters” (here, the selection based on the two independent subsets) with the chance agreement. Fisher exact test and *χ*^2^ association test quantify, respectively in a non-parametric and parametric way, the association between selections operated on the two subsets.

#### Classification accuracy

Classification accuracy was computed on a 10-fold cross-validation loop for *ProteoCardis* repeated 10 times to evaluate the standard deviation and a leave-one-out procedure leaving out all observations from each animal in turn on *Pigs*. The classification procedure consists in selecting the top *N* proteins, and then infer either a random forest (RF) or a support vector machine (SVM) based on these *N* features. For prediction, missing values were replaced by zero. Filtering was performed on the complete data set (as this step does not involve class labels). For each cross-validation split, the whole classification procedure including feature selection was performed on the training set only, then the labels of the samples in the validation set were predicted.

## RESULTS

### Missingness mechanisms: both MAR and MNAR

The replicate data sets, including seven technical replicates for eight biological samples, allow for the assessment of the technical variability and the analysis of the MAR/MNAR hypotheses. Figures 1 and S2 (left) display the average observed intensity of a protein as a function of the number of times it is missing in the replicate samples. The observed intensity decreases as the number of missing values increases, which suggests that the probability to be missing is higher when the protein abundance is lower, so missingness mechanisms is at least partially MNAR. In particular, we observed a pronounced difference of intensity between proteins with no missing value and protein with one, or *a fortiori* more than one, missing values. Nevertheless, even when the protein is missing in a large proportion of replicates, the average observed intensity can still be high, indicating that missingness mechanisms are not exclusively MNAR. These observations are confirmed by the probability of being missing, that decreases when the observed intensity increases, but remains non-negligible even for consistent observed intensities (Figures 1 and S2, right). For example, for an intensity equal to the median of the observed values on the data set, the probability of being missing is 0.23-0.25, and even for an intensity equal to the 0.9 quantile, the probability is still 0.05.

**Figure 1.**
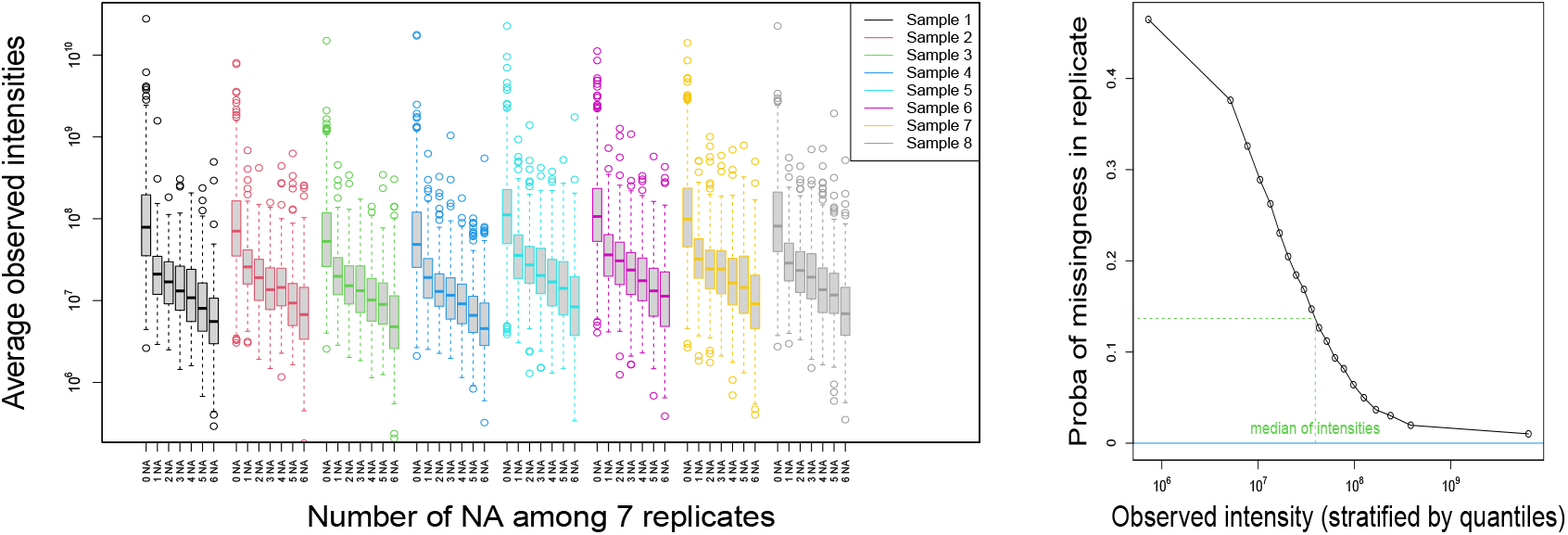
Analysis of replicates - cytosolic fraction. Left: log10-transformed average intensities of non-missing observations as a function of the number of missing values, for all proteins and for each biological sample. Right: Estimate of the probability that a protein is missing when the analysis is replicated, as a function of the average of its non-missing values.

### Comparison of FSM’s performances on simulated data

Figure 2 illustrates the ability to discriminate differentially and non-differentially expressed proteins of three FSMs, assuming two missingness mechanisms (MAR and MNAR) and under two types of difference of expression between proteins (differential presence and differential abundance). We considered two imputation-based FSMs: SVD based on global structure similarity, and single value imputation by the smallest observed intensity, followed by a linear (mixed) model, as well as the combined test. kNN-lmm was not considered since kNN imputation was used to generate the simulated data set. FSM’s performances are highly impacted by both the nature of the difference of expression and the missingness mechanism. Each imputation based FSM fails under one scenario: single-lmm does not achieve to detect differentially abundant proteins under MAR, while the AUC with SVD-lmm is close to 0.5 when proteins are differentially present under MNAR assumption. These observations are coherent with the underlying assumptions under each imputation methods: MAR for SVD and MNAR for single value imputation. In all scenarios, the combined test remains competitive, with AUCs close to the ones of the best of the two imputation-based FSMs.

**Figure 2.**
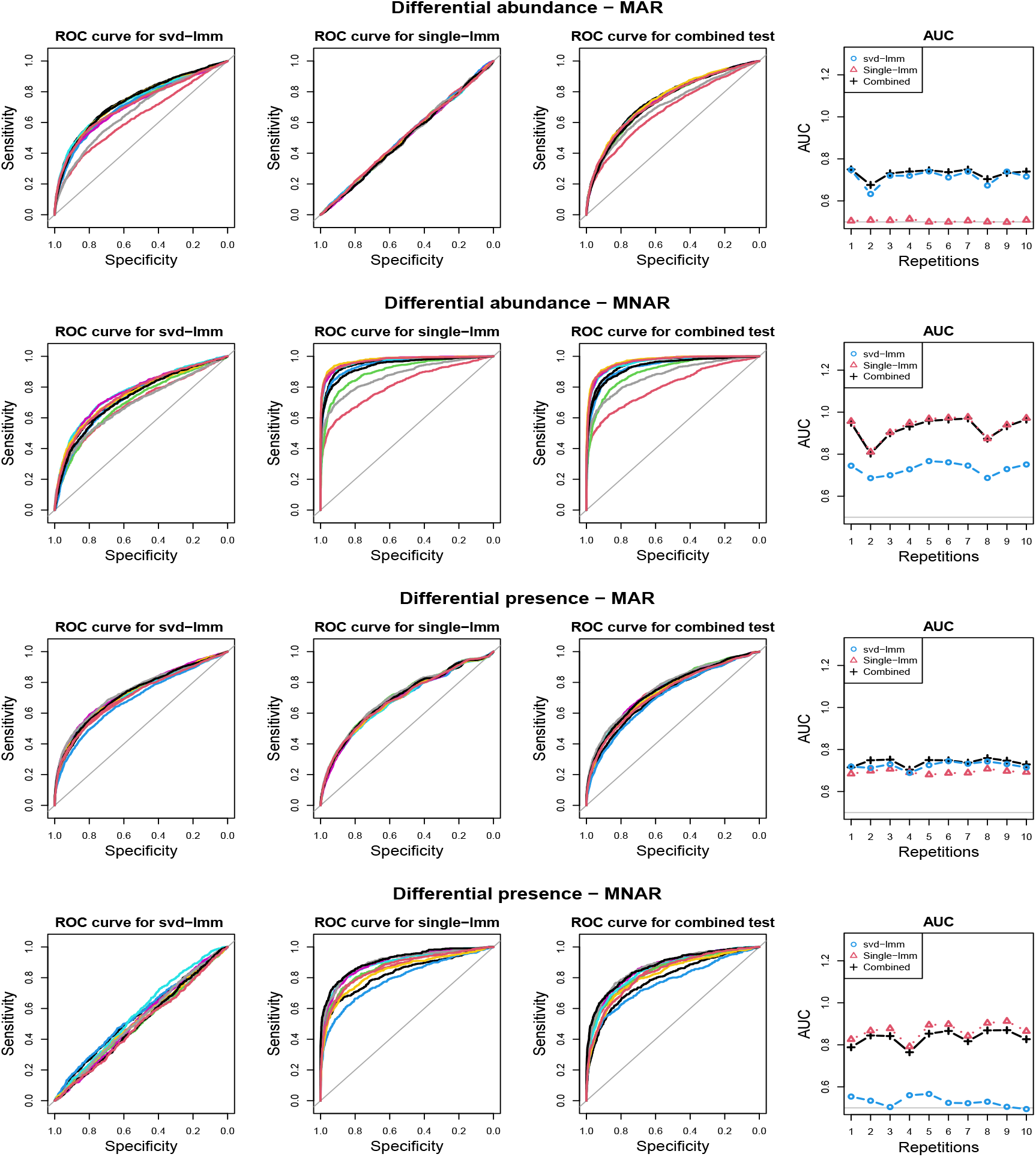
ROC curves for various scenarios on simulated data. Two types of difference between classes are generated: difference in intensity and difference in probability of absence, and two missingness mechanisms: Missing At Random and Missing Not At Random. For each scenario, the two classes and the proteins that differs between classes are randomly sampled 10 times, and the ROC curve with the three FSMs: linear model after SVD and single value imputation, and the combined test are displayed (each color correspond to a repetition. Column 4 displays the area under the curve for each method for the 10 repetitions.

### Poor concordance between imputation-based FSMs based on MAR and MNAR

Comparison of the three imputation-based FSMs in terms of correlation between log-transformed *p*-values and proportion of common selected features (Figures 3 and S3) indicates a strong agreement between SVD-lmm and KNN-lmm, but both methods show a poor concordance with single-lmm. These observations are common to the *ProteoCardis* and *Pigs* data sets, but the concordance between methods is globally higher for *Pigs* due to a weaker proportion of missing values, which reduces the impact of the imputation method.

**Figure 3.**
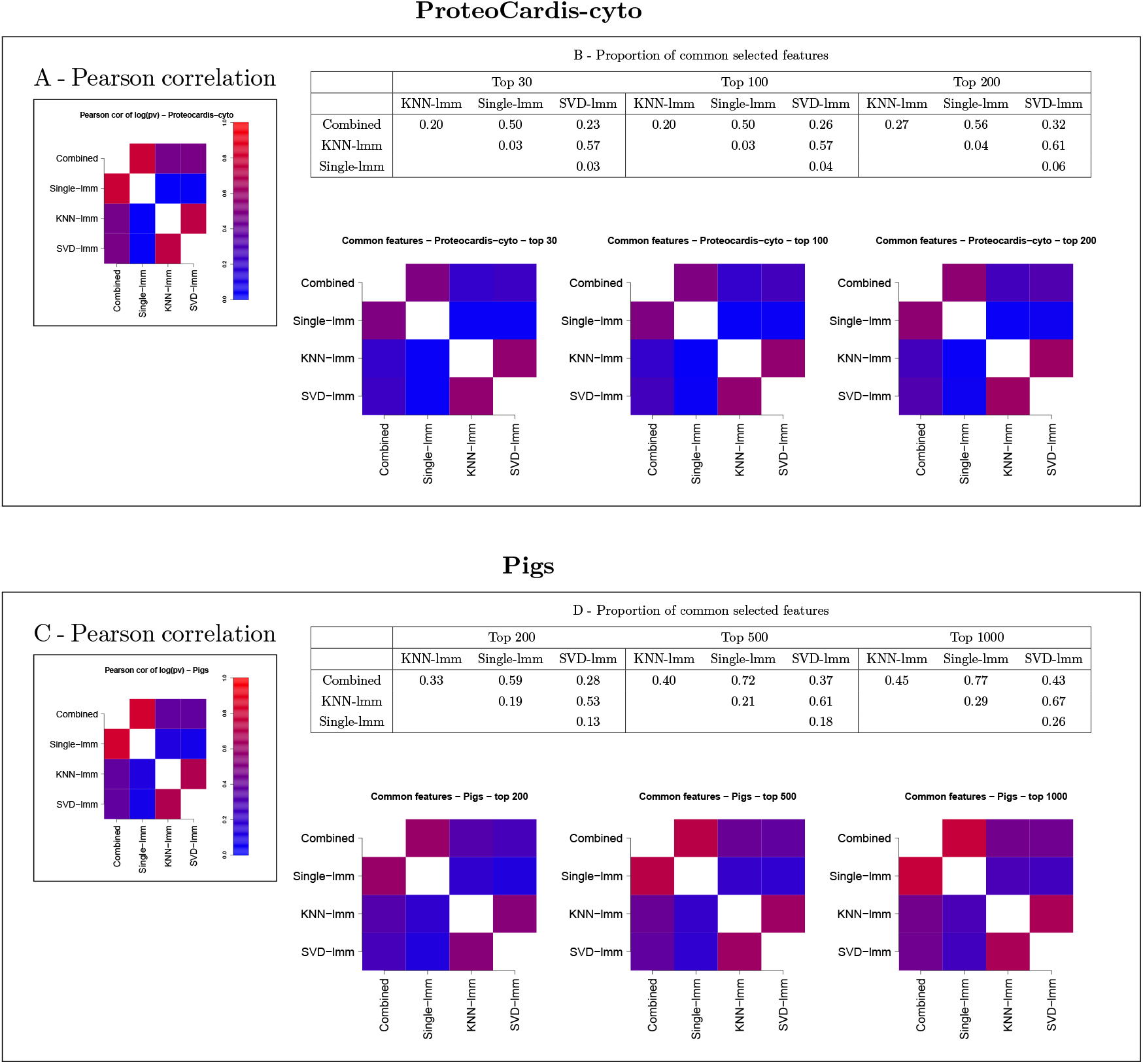
Pairwise agreement between *p*-values of FSMs. A,B: *ProteoCardis-cyto*; C,D: *Pigs*. A,C: Pearson correlation between log-transformed *p*-values. B,D: Proportion of common features among the top *N* for each pair of FSMs, as a table and a heatmap.

### Imputation-based FSMs are concordant with either glmm on probability of missingness or lmm on observed intensities

In addition, the imputation-based FSM *p*-values were compared with the two tests involved in the combined test (Rows 2 and 3 of Figures 4, 5 and S4). We observed a very strong correlation between the *p*-values of mod-glmm and single-lmm. Indeed, as the smallest intensity used for imputation is far from most of the observed intensities (Figure S9), the proportion of imputed values among a class strongly impacts the average intensity after single value imputation (a large proportion of missing values automatically leads to a small average intensity after single value imputation). Therefore, testing difference in fold-change and in missingness leads to consistent *p*-values. Single-lmm *p*-values are weakly consistent with mod-lmm on observed intensities. Conversely, the *p*-values from KNN-lmm and SVD-lmm correlate well with mod-lmm, but weakly with mod-glmm.

**Figure 4.**
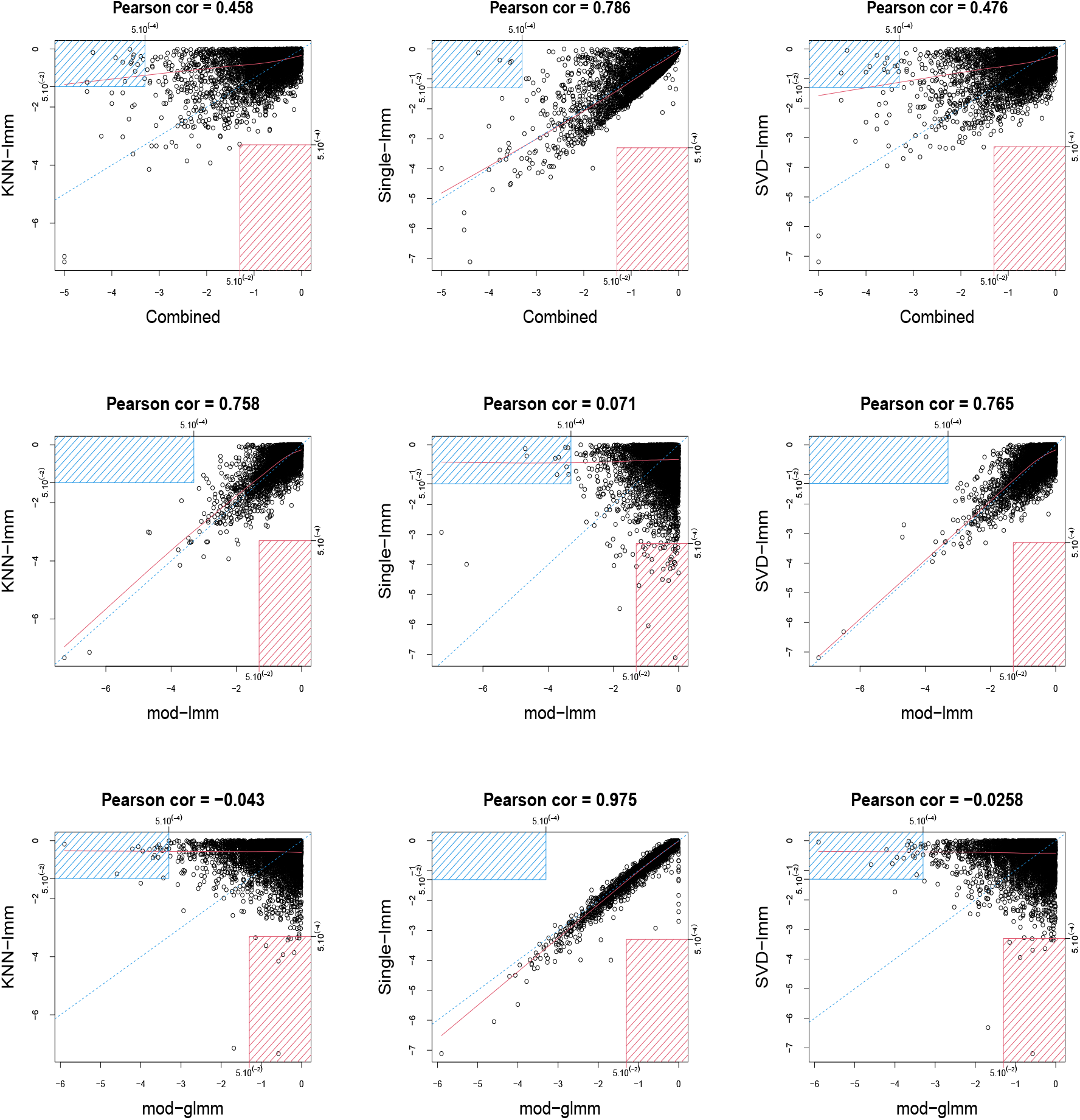
Scatterplots between log10-transformed *p*-values of pairs of FSMs for *ProteoCardis-cyto*. Row 1: combined test and imputation-based FSM. Row 2: Generalised mixed model (logistic) on missingness and imputation-based FSMs; proteins with less than 2 non-missing values are not displayed. Row 3: Linear mixed model on observed values and imputation-based FSMs. For each pair of testing procedure, the red rectangle corresponds to proteins with *p* > 5.10^-2^ with the first procedure and with *p* < 5.10^-4^ for the second procedure; conversely, the blue rectangle corresponds to proteins with *p* < 5.10^-4^ with the first procedure and with *p* > 5.10^-2^ for the second procedure.

**Figure 5.**
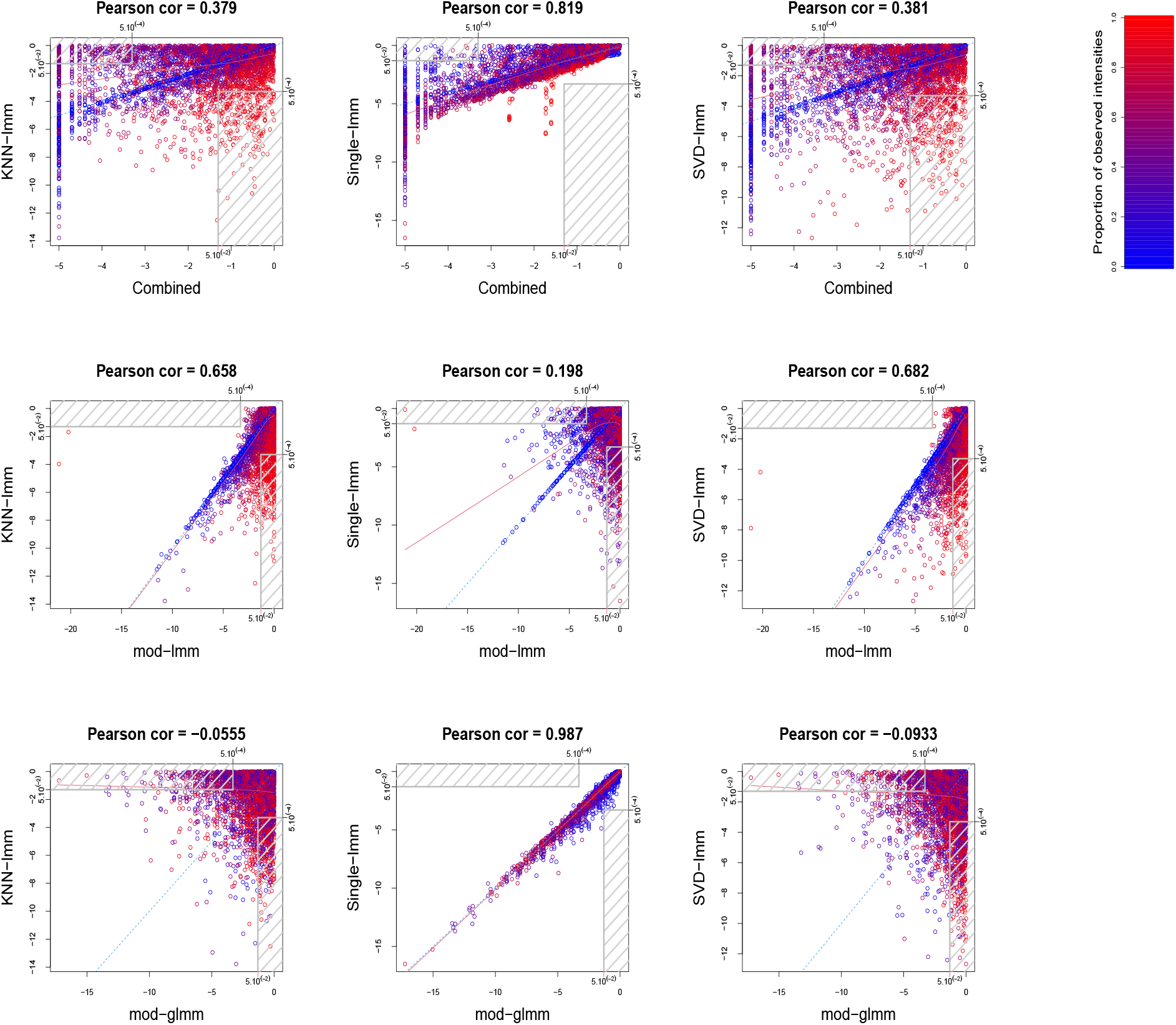
Scatterplots between log10-transformed *p*-values of pairs of FSMs for *Pigs*. Row 1: combined test and imputation-based FSMs. Row 2: Generalised mixed model (logistic) on missingness and imputation-based FSMs; proteins with less than 2 non-missing values are not displayed. Row 3: Linear mixed model on observed values and imputation-based FSM. Color gradient corresponds to the proportion of non-missing values for each protein. Gray rectangle correspond to features with *p* < 5.10^-4^ with one FSM and *p* > 5.10^-2^ with the other.

### The combined test reaches a compromise between imputation-based FSMs

The combined test displays a strong agreement with single-lmm and a moderate agreement with KNN-lmm and SVD-lmm in terms of correlation between log-transformed *p*-values and proportion of common selected features (Figures 3 and S3). This observation is confirmed by the scatterplots between log-transformed *p*-values of the combined test and each imputation-based FSM (first row of Figures 4, 5 and S4). Indeed, features found highly significant by any of the imputation based FSMs are at least moderately significant with the combined test, while the opposite is not true. More precisely, for *ProteoCardis* data sets, the proteins with very low *p*-values (*p* < 5.10 ^−4^) with the imputation-based FSMs also have low p-values with the combined test (*p* < 5.10^-2^); Conversely for each imputation-based FSM, some non-significant proteins display very low *p*-values with the combined test. A more nuanced but similar assessment holds on *Pigs*, since most of the proteins that are found significant with SVD-lmm or KNN-lmm and non-significant with the combined test are very sparse (Figure S5), and thus include a large proportion of imputed values, which indicates that imputation-based analysis on these variables is weakly reliable. On the contrary, the variables significant with the combined test but non-significant with SVD-lmm and KNN-lmm have a sparsity level that is either low or high.

### Impact of the filtering threshold

The impact of the imputation method is expected to decrease with the proportion of missing values, which itself depends on the filtering threshold. Therefore, we repeated our analyses with higher filtering thresholds: 30, 40 and 50 for *ProteoCardis*, and 20 and 30 for *Pigs*, to examine to what extent the comparisons between FSMs were impacted. Figures S6, S7 and S8 display the concordance between FSMs for several filtering thresholds. The comparisons between methods still holds when threshold varies: SVD-lmm/KNN-lmm on the one hand, and single-lmm/combined test on the other hand are strongly concordant, while KNN-lmm/SVD-lmm are moderately concordant with the combined test, and poorly concordant with single-lmm. As expected, the agreement between all pairs of FSMs globally increases with the threshold.

A harder thresholding also results in a higher number of discarded variables, potentially leading to a loss of biological information. Indeed, Figure S10 displays the distribution of the *p*-values as a function of the protein sparsity; we observed that very sparse proteins may exhibit very small *p*-values, in particular for *ProteoCardis* data sets. Moreoever, Table S1 indicates that the sets of most significant proteins for each FSM include 60-82% (average 70%) of features with more than half of missing values for *ProteoCardis*, and 17-57% (average 41%) for *Pigs*. Note that this proportion of sparse proteins among the selected ones is only slightly lower than in the entire data sets (79% for *ProteoCardis* and 43% for *Pigs*). This indicates that sparse proteins are selected almost as frequently as less sparse ones.

### Similar FSM’s quantitative performances

Tables 1 and S2 display the prediction accuracy with SVM and RF classifiers applied on selected proteins. Even though the four FSMs select very different sets of features, their performances in terms of prediction are not significantly different.

**Table 1.**
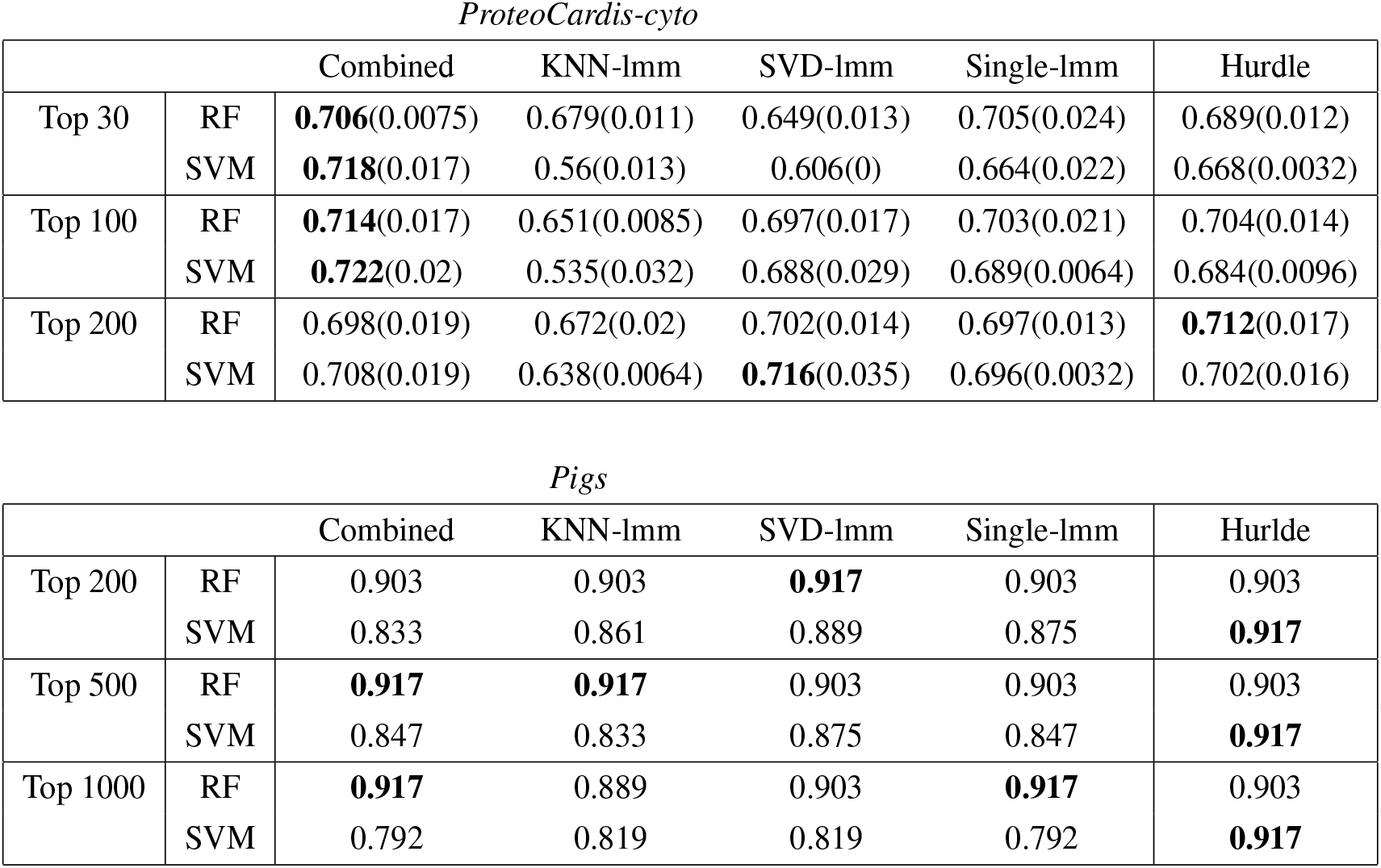
**Prediction accuracy** for two classification procedures on *ProteoCardis-cyto* and *Pigs*. The selection of the top *N* variables was followed by SVM or RF. For *Proteocardis-cyto*, accuracy was computed in a 10-fold cross-validation loop, repeated 10 times. For *Pigs*, accuracy was computed in a leave-one-out setting in which training sets consist in all measurements from one pig. Each cell provides the average accuracy (standard deviation of accuracy) computed over the 10 repetitions of the cross-validation. Bold numbers correspond to the highest accuracy among the four FSMs.

*ProteoCardis-cyto*
|  |  | Combined | KNN-lmm | SVD-lmm | Single-lmm | Hurdle |
| --- | --- | --- | --- | --- | --- | --- |
| Top 30 | RF | **0.706**(0.0075) | 0.679(0.011) | 0.649(0.013) | 0.705(0.024) | 0.689(0.012) |
|  | SVM | **0.718**(0.017) | 0.56(0.013) | 0.606(0) | 0.664(0.022) | 0.668(0.0032) |
| Top 100 | RF | **0.714**(0.017) | 0.651(0.0085) | 0.697(0.017) | 0.703(0.021) | 0.704(0.014) |
|  | SVM | **0.722**(0.02) | 0.535(0.032) | 0.688(0.029) | 0.689(0.0064) | 0.684(0.0096) |
| Top 200 | RF | 0.698(0.019) | 0.672(0.02) | 0.702(0.014) | 0.697(0.013) | **0.712**(0.017) |
|  | SVM | 0.708(0.019) | 0.638(0.0064) | **0.716**(0.035) | 0.696(0.0032) | 0.702(0.016) |

**Table 1. Prediction accuracy** for two classification procedures on *ProteoCardis-cyto* and *Pigs*.
|  |  | Combined | KNN-lmm | SVD-lmm | Single-lmm | Hurdle |
| --- | --- | --- | --- | --- | --- | --- |
| Top 200 | RF | 0.903 | 0.903 | **0.917** | 0.903 | 0.903 |
|  | SVM | 0.833 | 0.861 | 0.889 | 0.875 | **0.917** |
| Top 500 | RF | **0.917** | **0.917** | 0.903 | 0.903 | 0.903 |
|  | SVM | 0.847 | 0.833 | 0.875 | 0.847 | **0.917** |
| Top 1000 | RF | **0.917** | 0.889 | 0.903 | **0.917** | 0.903 |
|  | SVM | 0.792 | 0.819 | 0.819 | 0.792 | **0.917** |

Figures 6 and S11 display the concordance between variable selection performed on independent data sets. On *ProteoCardis-cyto*, reproducibility of feature selection is similar with the four FSMs, while on *ProteoCardis-env* and *Pigs* the combined test and single-lmm outperform imputation-based FSMs. Therefore variable selection based on the combined test is equally reproducible that single imputation FSM, and tends to be more reproducible than FSMs using structure-based imputation.

**Figure 6.**
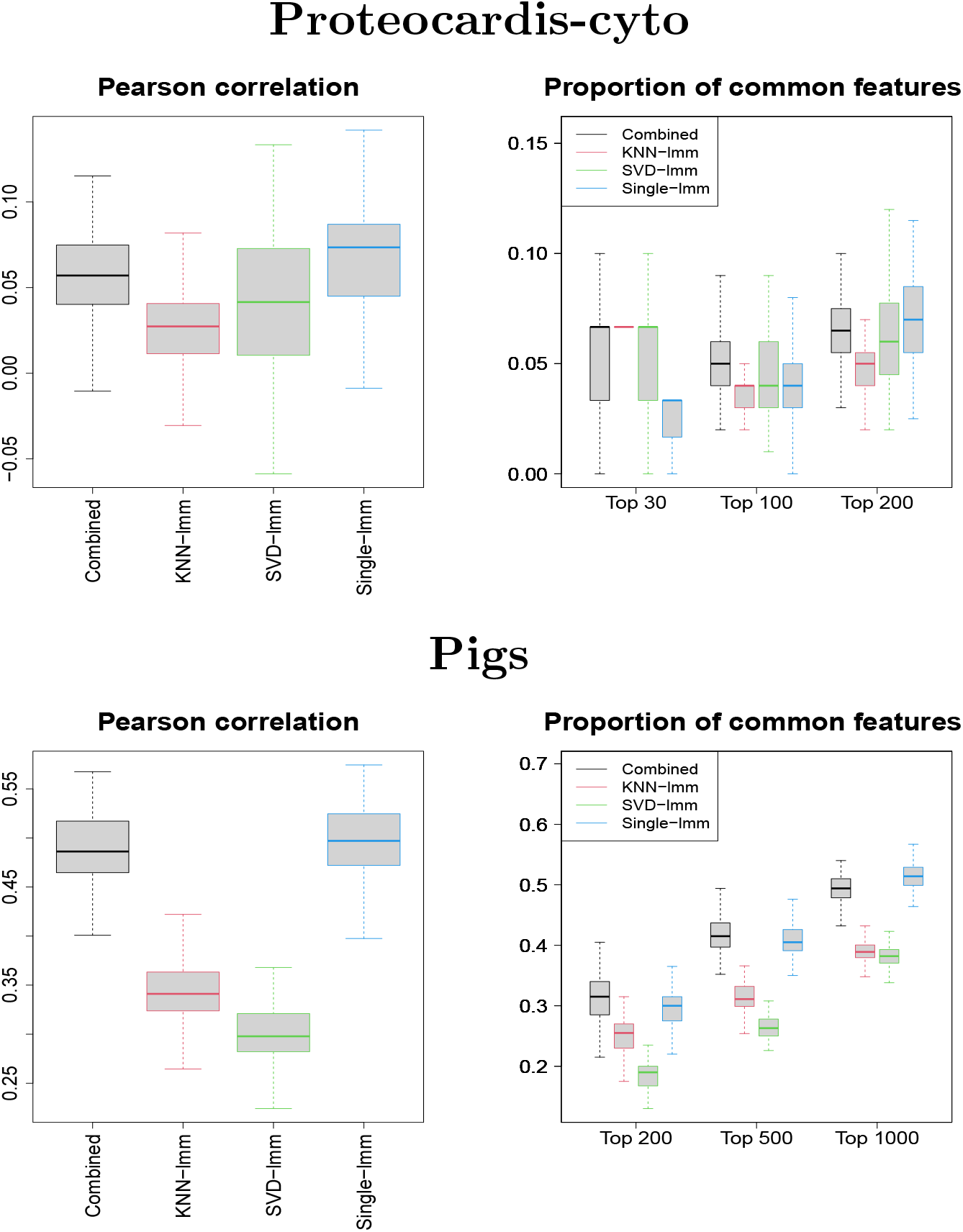
Replicability of variable selection on independent subsets. Pearson correlation between log10-transformed *p*-values and proportion of common selected features among the top *N*, for 100 splitting of samples into two subsets. Datasets: *ProteoCardis-cyto* and *Pigs*

Figure S1 displays the number of selected variables for various values of FDR. The methods ranking varied with the data set and the FDR threshold, but the combined test remains competitive in terms of number of selected variables.

### Comparison of the combined test with the peptide-level hurdle model

Similarly to the combined test, the Hurdle model proposed by Goeminne et al. (2020) targets simultaneously difference of intensity on observed values and difference in probability of detection, but with a peptide level model. As expected, both procedures lead to consistent *p*-values (Figure 7), with Pearson correlation of log-transformed *p*-values of 0.637-0.786, but far from identical. In terms of feature selection, some proteins highly significant with one method exhibit non-significant *p*-values with the other, in particular for *Pigs*.

**Figure 7.**
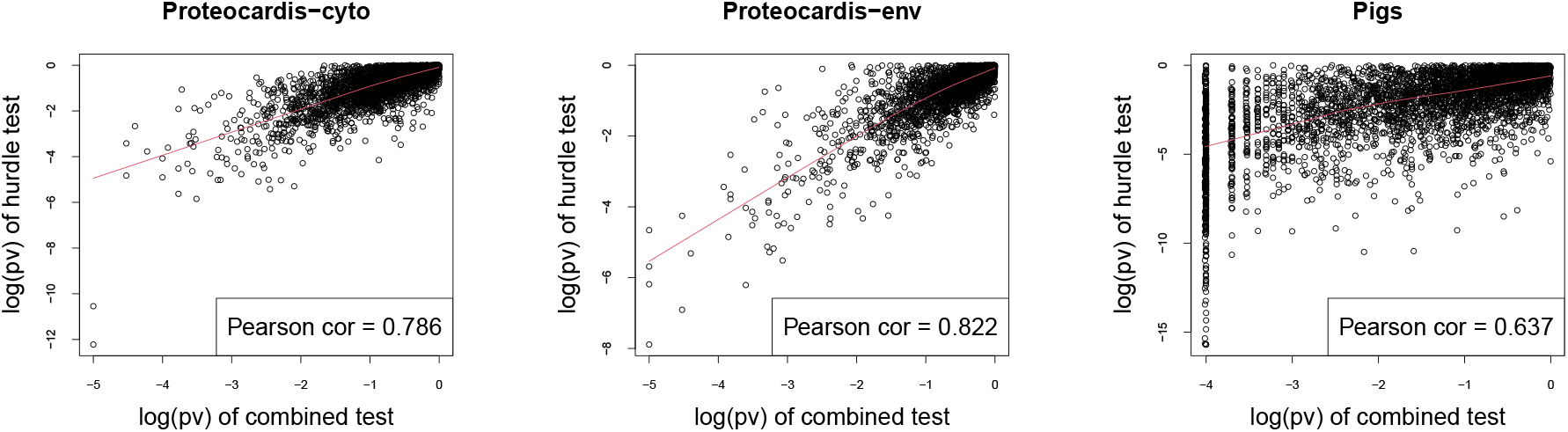
Scatterplot of log10-transformed *p*-values of the hurdle test and the combined test for *ProteoCardis-cyto* and *Pigs* data sets.

Regarding prediction, both procedures lead to similar accuracy on *ProteoCardis* data sets, and on *Pigs* with RF, but the combined test has less good performances with SVM than RF, while the hurdle test maintain a similar accuracy (Tables 1 and S2). No significant differences in terms of replicability were observed between the protein-level combined test and the peptide-level hurdle test (Figures 8 and S12). More precisely, feature selection with the combined test is slightly more replicable than the hurdle test on *Pigs*, notably for the largest number of selected features, while the opposite is observed on *ProteoCardis* data sets.

**Figure 8.**
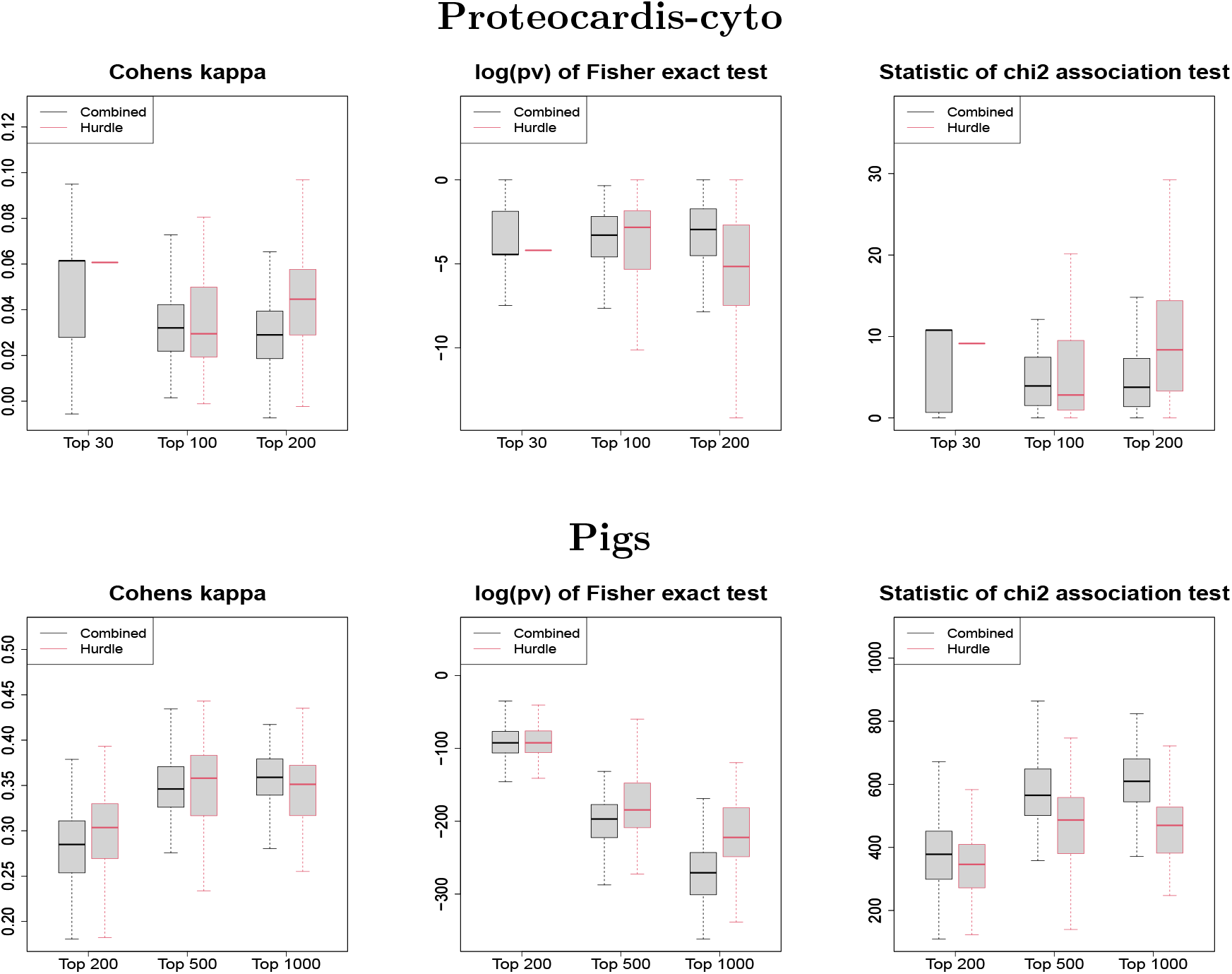
Replicability of variable selection on independent subsets for the hurdle test and the combined test. Boxplot of the Cohen’s kappa (left), the log-transformed *p*-value of Fisher test (center) and the statistic of the *χ*^2^ contingency table test (right), for selection of the top *N* features, performed on 100 splitting of the samples into two subsets. Black and red boxlots correspond to feature selection with the combined and the hurdle test respectively. Datasets: *ProteoCardis-cyto* and *Pigs*

## DISCUSSION

### Missingness blends MAR and MNAR mechanisms and our method addresses both assumptions

Metaproteomics by LC-MS/MS generates a large proportion of missing values, usually imputed prior to statistical analysis. Several categories of imputation methods are routinely considered. Methods based on local similarity (e.g. kNN) or global structure (e.g. SVD) implicitely assume that missingness occurs independently of the true feature concentration (MAR). But the analysis of the replicate data sets clearly indicates that missingness is more likely to occur when the feature has a low abundance. On the other hand, the single imputation method relies on the assumption of a left censoring mechanism, but the distribution of observed intensities as well as the analysis of the replicate data set are not consistent with this assumption. Even if the co-existence of MAR and MNAR mechanisms is admitted in LC-MS/MS proteomics, exploration of their prevalence is often based on biological replicates (e.g. Karpievitch et al. (2009)), assuming similar protein abundances in all samples from a given class. This assumption is questionable, in particular in human gut metaproteomics characterised by a strong individual specificity. On the contrary, our analysis based on technical replicates guarantees that the true protein abundances are identical.

### Limits of missing value imputation in metaproteomics

Missing value imputation is the standard method to account for missing values in metaproteomics. This flexible approach enables to address any type of statistical questions (e.g. prediction, network inference…) using methods developed for data with no missing values. But the downside is the risk to “forget” which values were imputed and to treat them equally to observed values, regardless of implicit assumptions underlying imputation that can strongly impact biological findings when a large proportion of values are missing (O’Brien et al., 2018; Karpievitch et al., 2012; Lazar et al., 2016). In particular, we observed that global and local structure imputations specifically led to selection of features with a large proportion of imputed and thus weakly reliable values. Therefore, despite its easiness of use, imputation has a cost in terms of reliability and should be limited to moderately sparse data sets. On sparse metaproteomics data, this condition would require to filter out a large part of the features, which may be harmful since we demonstrated that a large part of the potentially interesting proteins have more than half of missing values.

As an alternative to missing value imputation, censored statistical models developed for proteomics data can account simultaneously for MAR and MNAR mechanisms (Karpievitch et al., 2009; Luo et al., 2009; O’Brien et al., 2018). Moreover, Berg et al. (2019) proposed a multiple imputation model which handles MAR and MNAR assumptions, but their method suffers from a methodological bias since imputation is performed within each class, which may artificially increase significance of inter-classes differences by enhancing intra-class similarities. Similarly, Gianetto et al. (2020) proposed a combination of MAR and MNAR imputation for proteomics data, but their method relies on a log-normal distribution of the true abundances, unrealistic in the metaproteomics framework where a large part of proteins are truly missing. Even though these models may address the complexity of missingness mechanisms more acutely than simple imputation methods, they also heavily rely on assumptions regarding missingness mechanisms (e.g. hard thresholding) as well as signal distribution (e.g. additive effect, Gaussian distribution). Therefore, they can not be directly applied to metaproteomics data whose structure and characteristics strongly differ from proteomics.

### The combined test reaches a compromise between imputation-based methods

Beyond differences in underlying assumptions, distinct imputation-based FSMs lead to very different sets of selected variables. While selection with the two MAR imputation methods (SVD and kNN) are consistent, these two methods display almost no agreement with the FSM based on MNAR assumption (single value imputation) for the highly sparse data sets *ProteoCardis*, and a low agreement for the moderately sparse data set *Pigs*. Therefore, the choice of an imputation method can strongly impact the biological conclusions. The combined test, which addresses the two types of missingness by combining a glmm on probability of presence, which is relevant under MNAR assumption, and a lmm on observed intensities after removal of missing values, which amounts to assume MAR mechanisms displays a correct agreement with each imputation-based FSM. In greater details, the features detected using single value imputation were recovered by the glmm, while the features selected using kNN or SVD were recovered by the lmm on observed values, which is consistent with the assumptions on missingness mechanisms.

Besides, prediction accuracy is a classic criterion to compare FSMs (Tang et al., 2020b), but its relevance is questionable when the main interest is the biological interpretation of selected features. Indeed, as enlightened in our analysis, methods with similar classification abilities can lead to totally different selected sets and feature ranking, so choosing a FSM based on a slightly higher prediction accuracy - that may vary with the chosen classifier - seems hazardous.

Furthermore, the simulation study conducted on archetypical scenarios illustrates that the comparative performances in terms of discrimination between differentially and non-differentially expressed proteins strongly depends on the nature of the protein difference (associated to the biological context) and the missingness mechanisms (mostly dependent on the technology). First of all, this simulation study indicates that quantitative comparison of FSMs’ performances in a specific context has to be taken cautiously in another biological context, in which the proportion of each scenario, notably regarding biological differences, could vary. In particular, transposition of methods comparisons performed on proteomics data to the metaproteomics context can be hazardous, as both the biological context and the technical artefacts can differ. Secondly, while each imputation-based FSM fails in at least one scenario, the combined remains efficient in all scenarios.

Thus, the combined test realises a compromise in terms of feature ranking and selection between inconsistent methods whose performances are based on unverifiable assumptions, and remains efficient in diverse scenarios. Therefore, it represents a robust solution, notably in the context of few prior knowledge regarding missingness mechanisms and the nature of biological differences.

### Protein versus peptide level analysis

In proteomics, analyses can either be realised at the protein or peptide level. In univariate testing procedures on proteomics data, peptide level analyses including a run effect are usually considered as more robust and with higher power (Clough et al., 2012). Nevertheless, in metaproteomics, analysis are routinely performed at the protein level (see review by Tang et al. (2020b)). This choice enables simplifications in the analysis, but biological arguments could be considered: the complexity of metaproteome may lead to a higher proportion of peptide misidentifications; moreover, the individual specificity of the metaproteome generates a very high sparsity at the peptide level (e.g. 95-97% of missing values at the peptide level in Proteocardis data sets) which is detrimental to the robustness of mixed model.

We compared the combined test with the hurdle model by Goeminne et al. (2020), which displays similarities but is implemented at the peptide level. Interestingly, contrary to what was demonstrated by the authors on proteomics data, this peptide level analysis did not demonstrate superior performances compared to the protein-level combined test on our metaproteomics data sets. Further analyses of the biological findings brought by each method would enrich the comparison.

Beyond metaproteomics, the generic aspect of the combined test enables to use it in other omics or non-omics contexts with data missing both at random and not at random.

## CONCLUSION AND PERSPECTIVES

Feature selection based on imputation is highly dependent on the chosen imputation method, and thus on restrictive assumptions regarding missingness mechanisms, while biometric measurements are most of the time subject to mixed missingness processes. Moreover, beyond censoring mechanisms, we enhanced that FSMs’ performance are strongly impacted by the type of difference of expression. On the contrary, the combined test handles simultaneously missingness at random and not at random, and our analysis on metaproteomics data confirm its effectiveness to recover the strongest findings from imputation-based FSM based on either type of mechanisms and for different nature of biological changes, while displaying equivalent quantitative performances.

In this paper, we focused on the missing data issue and we restricted our analysis to FSMs based on a linear mixed model, but the combined test could further be compared with more diverse FSMs, including wrapped and embedded methods (Tang et al., 2020b), as well as using data sets whose design include more than two classes. On the biological side, the conclusion of this article could be reinforced through validation by targeted proteomics measurements on a subset of variables. Ground truth data sets such as spike-in, where the concentration of a small number of features is controlled could also be considered, but one should keep in mind that comparative analyses of FSMs are strongly impacted by the type of biological differences between classes (notably differential presence or abundance). Finally, the combined test developed in this article is not restricted to metaproteomics data and could be implemented on other meta-omics data or on any data including a large part of missing values, whatever the missingness mechanisms. Moreover, the proposed approach could be generalised to univariate feature selection in other frameworks than multi-class comparison (e.g. time series) provided that a test of presence/absence is available (e.g. a rank test for time series).

## Acronyms

FSM: Feature Selection Method
FDR: False Discovery Rate
kNN: k-Nearest Neighbors
SVD: Singular Value Decomposition
SVM: Support Vector Machine
RF: Random Forest
MAR: Missing At Random
MNAR: Missing Not At Random

## CODE AND DATA AVAILABILITY

Software in the form of R code, together with the ProteoCardis data sets are available at https://doi.org/10.15454/ZSREJA.

## CONTRIBUTION STATEMENTS

S. Plancade and M. Berland conceived the original idea, performed the numerical implementation, analysed the data and wrote the article. M. Blein Nicolas and C. Juste participated to discussions about the method and revised critically the draft. ProteoCardis data were produced by C. Juste and pre-processed by O. Langella and A. Bassignani from samples provided by the Metacardis European FP7 initiative, and prepared and analyzed for the ANR project Proteocardis.

## ACKNOWLEDGMENTS

This work was supported by the Agence Nationale de la Recherche (ANR) as part of the ProteoCardis (ANR-15-CE14-0013) project. The proteomics analyses were performed on the PAPPSO facility (http://pappso.inrae.fr) which is supported by INRAE (http://www.inrae.fr), the Ile-de-France regional council (https://www.iledefrance.fr/education-recherche), IBiSA (https://www.ibisa.net) and CNRS (http://www.cnrs.fr). We thank Alfred AMEADAN (Univ. Paris-Saclay, INRAE, APPSO-Micalis, 78350, Jouy-en-Josas, France) for preparing, analysing, and generating the raw LC-MS/MS data sets of the ProteoCardis study. Sylvie HUET (senior research scientist at MaIAGE-INRAE, UniversitÃ© Paris-Saclay, 78350 JOUY-en-JOSAS), Christine CARAPITO (senior research scientist at IPHC-CNRS, 67037 STRASBOURG) and Magali ROMPAIS (research engineer at LSMBO, IPHC-CNRS, 67037 STRASBOURG) are gratefully acknowledged for their support and valuable contribution to scientific discussions.

## SUPPLEMENTARY MATERIALS

### Permutation procedure for the combined test

#### Preliminary: define the permutation design

According to the study design, the user can provide constraints on the permutations *via* control parameters defined by the function `how` to be passed to the R function `shuffle` (`permute` package). For the *ProteoCardis* datasets, no constraints were considered, but for *Pigs*, classes were permuted while keeping together the samples from the same animal.

#### Permutation test p-values

Let *n* be the number of biological samples in the dataset, and *m* the number of proteins. Let (*a_j_*)_*j*=1,…,*m*_ be the number of non-missing intensities among the *n* samples for each protein *j* = 1,…, *m*. After filtering of proteins with less than *τ* non-missing values, *a_j_* ∈ {*τ*,…, *n*} for all *j*. Then, for each *a* ∈ {*τ*,…, *n*},

- Let 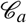 be the set of proteins with *a* non-missing values.
- For each protein 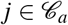, classes are permuted repeatedly according to the chosen permutation design, for repetitions 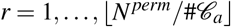 with ⎣·⎦ the integer part and # the cardinal, and the Fisher combined statistic 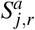 is computed.
- The vector 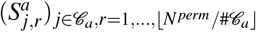 represents a sample of the distribution of the test statistic under the null hypothesis of no class effect, for proteins with *a* non-missing values.

Then, for each protein *j* = 1,…, *m*, the *p*-value of the combined test is equal to:

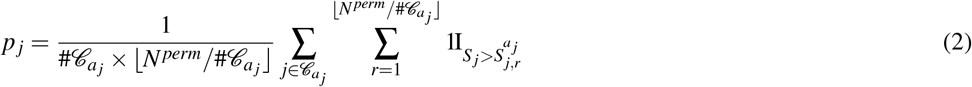

with *S^j^* the Fisher combined statistics of protein *j* computed with the true classes.

#### Resampling based FDR

Resampling-based FDR is computed for 100 permutations. For *s* = 1,…, 100,

- Classes are permuted according to the chosen permutation design.
- The Fisher combined statistic 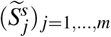 is computed, using the same permuted classes for all proteins.
- The vector 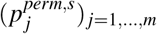 of *p*-values under the complete null assumptions are computed by equation (2) with *S_j_* replaced by 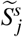. Note that the distribution under the null assumption does not require to be computed again.

Following the procedure by Reiner et al. (2003), new estimates of the *p*-values are computed assuming that the marginal distributions under the complete null hypothesis are exchangeable:

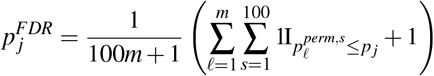

Finally, FDR adjustment (Benjamini and Hochberg, 1995b) is applied to 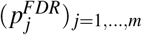.

### Simulation framework

#### General procedure

- Protein intensities from the data set *ProteoCardis-cyto* were filtered at threshold 10 (i.e. proteins with less than 10 non-missing values were removed), resulting in 11,433 proteins and 74% of missing values. Then the missing values were imputed by kNN, providing a realistic metaproteomic data set.
- Two classes of size 49 and 50 were randomly sampled among the 99 samples.
- 2000 proteins were randomly selected to be different between the two classes. Two types of difference were considered: (i) Differential intensity, (ii) Differential presence (see details below).
- Two missingness scenarios were considered: (ii) MAR: Missing values were drawn randomly such that the proportion of missing values on the total data set is equal to the proportion in the original data set *Proteocardis-cyto* after filtering at level 10; (ii) MNAR: a hard thresholding was applied, with threshold chosen to have the same proportion of missing values than on *ProteoCardis-cyto* after filtering at level 10.
- For the 2 × 2 scenarios, proteins with less than 20 non-missing values were removed, then the three FSMs SVD-lmm, single-lmm and the combined test were implemented, and the ROC curves were computed. Note that KNN-lmm was not considered since it includes the same imputation method used to generate the data set; Besides, this method has been shown to perform similarly to SVD-lmm.

#### Generate difference between groups

- **Differential intensity**. For each of the 2000 proteins, the quantity *FC_j_*/2 was added to the intensities of samples from one class and substracted to the intensities of samples from the other class. The fold change *FC_j_* was tuned so that the corresponding *p*-value of a t-test was approximately equal to *α* = 10 ^−3^, according to the standard deviation σ_j_ of the intensities of each protein. More precisely, for a fold-change *FC_j_*, the t-test statistic is equal to

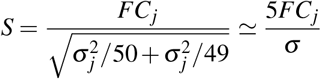 Thus, setting the *p*-value to *α* is equivalent to:

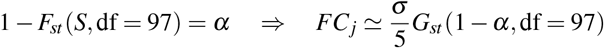

where *F_st_* and *G_st_* denote the cumulative distribution function and the quantile function of the student distribution.
- **Differential presence.** For each of the 2000 proteins, each intensity was set to NA with probability *τ* in one class and 1 – *τ* in the other class. The parameter *τ* was tuned such that the *p*-value of the Fisher exact test for the average table:

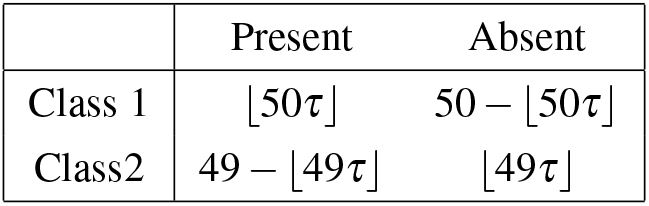

was equal to *α* = 10 ^3^, where ⎣·⎦ denotes the integer part.

## SUPPLEMENTARY FIGURES AND TABLES

**Table S1.** **Proportion of selected variables with less than half observed intensities**, among the top N variables (between 20 and 50 non-missing values for *Proteocardis* data sets, and between 10 and 36 for *Pigs*).

|  | Proteocardis-cyto |  |  | Proteocardis-env |  |  | Pigs |  |  |
| --- | --- | --- | --- | --- | --- | --- | --- | --- | --- |
|  | top30 | top100 | top200 | top30 | top100 | top200 | top200 | top500 | top1000 |
| Combined | 0.60 | 0.69 | 0.69 | 0.63 | 0.68 | 0.76 | 0.17 | 0.17 | 0.25 |
| KNN-lmm | 0.67 | 0.63 | 0.68 | 0.70 | 0.76 | 0.82 | 0.57 | 0.57 | 0.57 |
| SVD-lmm | 0.70 | 0.65 | 0.69 | 0.67 | 0.74 | 0.80 | 0.28 | 0.31 | 0.34 |
| Single-lmm | 0.60 | 0.68 | 0.72 | 0.60 | 0.69 | 0.78 | 0.60 | 0.57 | 0.54 |

**Figure S1.**
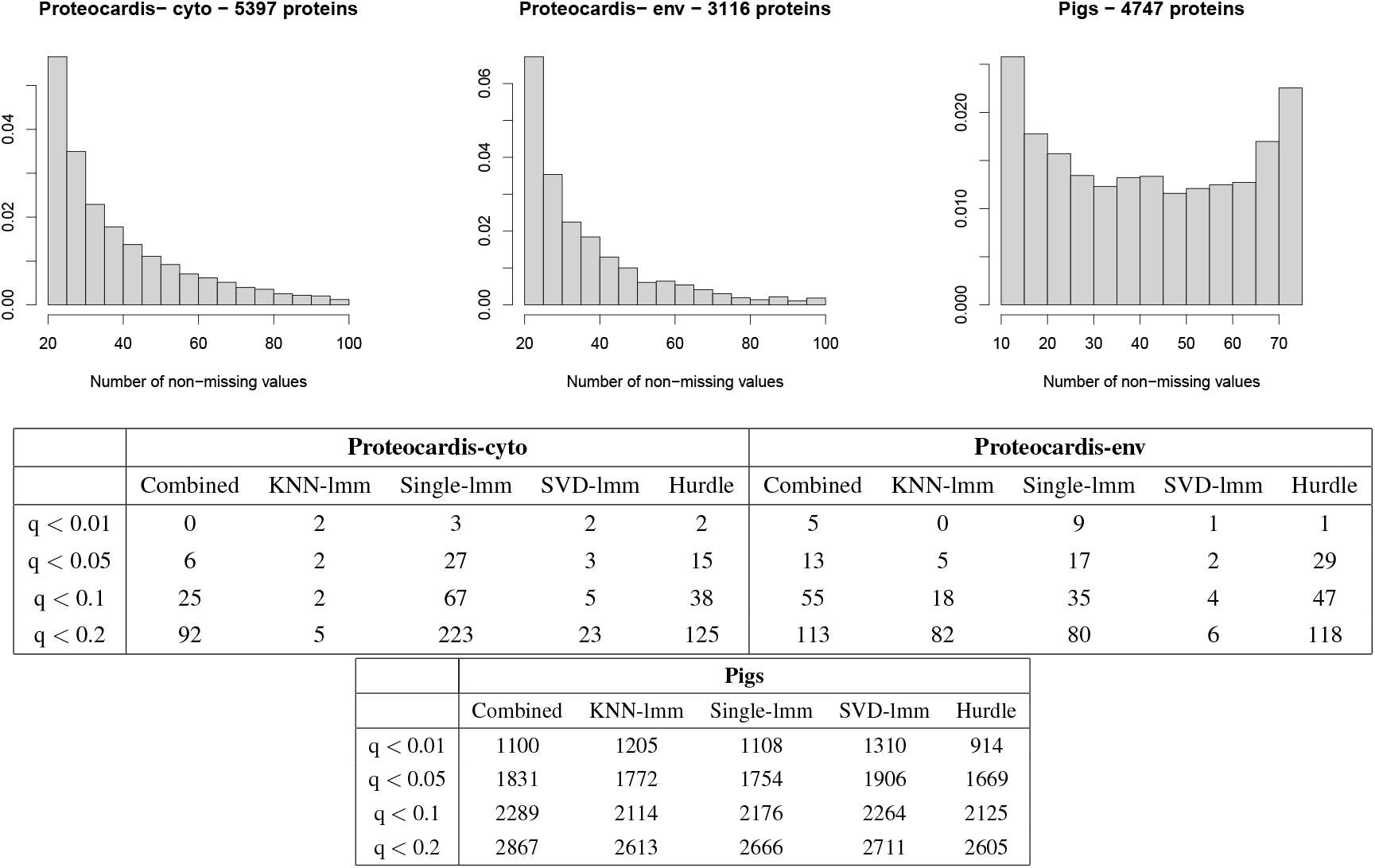
Statistical characteristics of the three data sets *ProteoCardis-cyt*, *Proteocardis-env*, *Pigs*. Top: frequencies of the number of non-missing values for all proteins after filtering (threshold 20 for *ProteoCardis*, and 10 for *Pigs*). Bottom: number of selected variables with the resampling FDR procedure with 100 resampling repetitions, with various values of the FDR threshold values.

**Figure S2.**
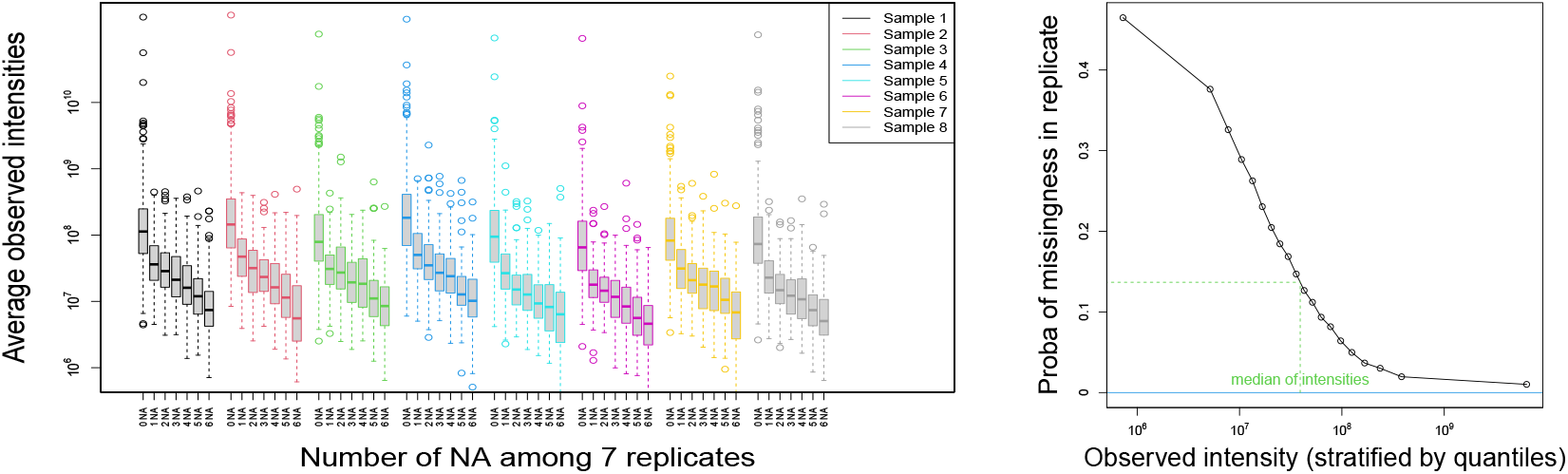
Analysis of replicates - envelope fraction. Left: log10-transformed average intensities of non-missing observations, as a function of the number of missing values, for all proteins and for each biological sample. Right: Estimate of the probability that a protein is missing in a technical replicate as a function of the average of its non-missing values.

**Figure S3.**
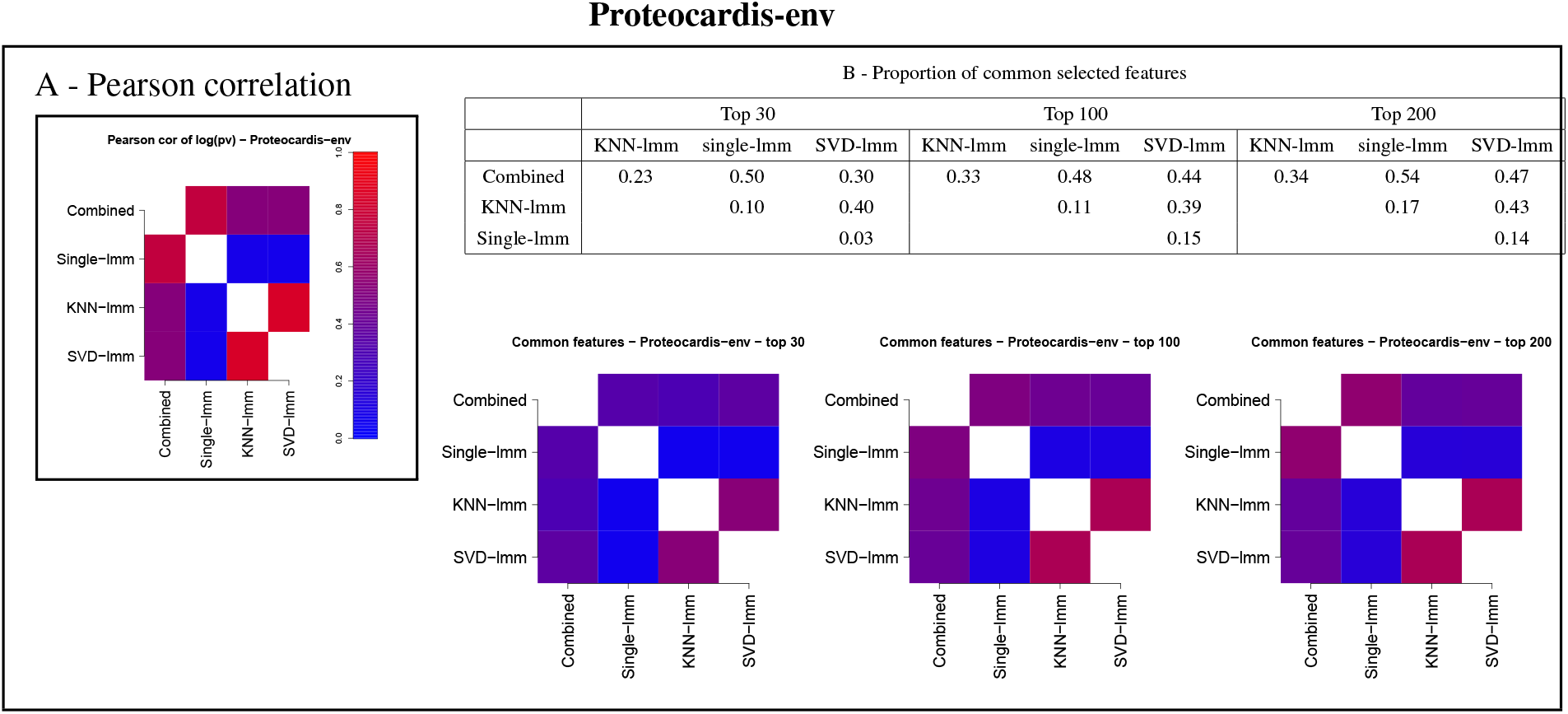
Pairwise agreement between *p*-values of FSMs for *Proteocardis-env*. A: Pearson correlation between log of *p*-values. B: Proportion of common features among the top *N* (*N* = 30, 100, 200) for each pair of FSMs, as a table and a heatmap.

**Figure S4.**
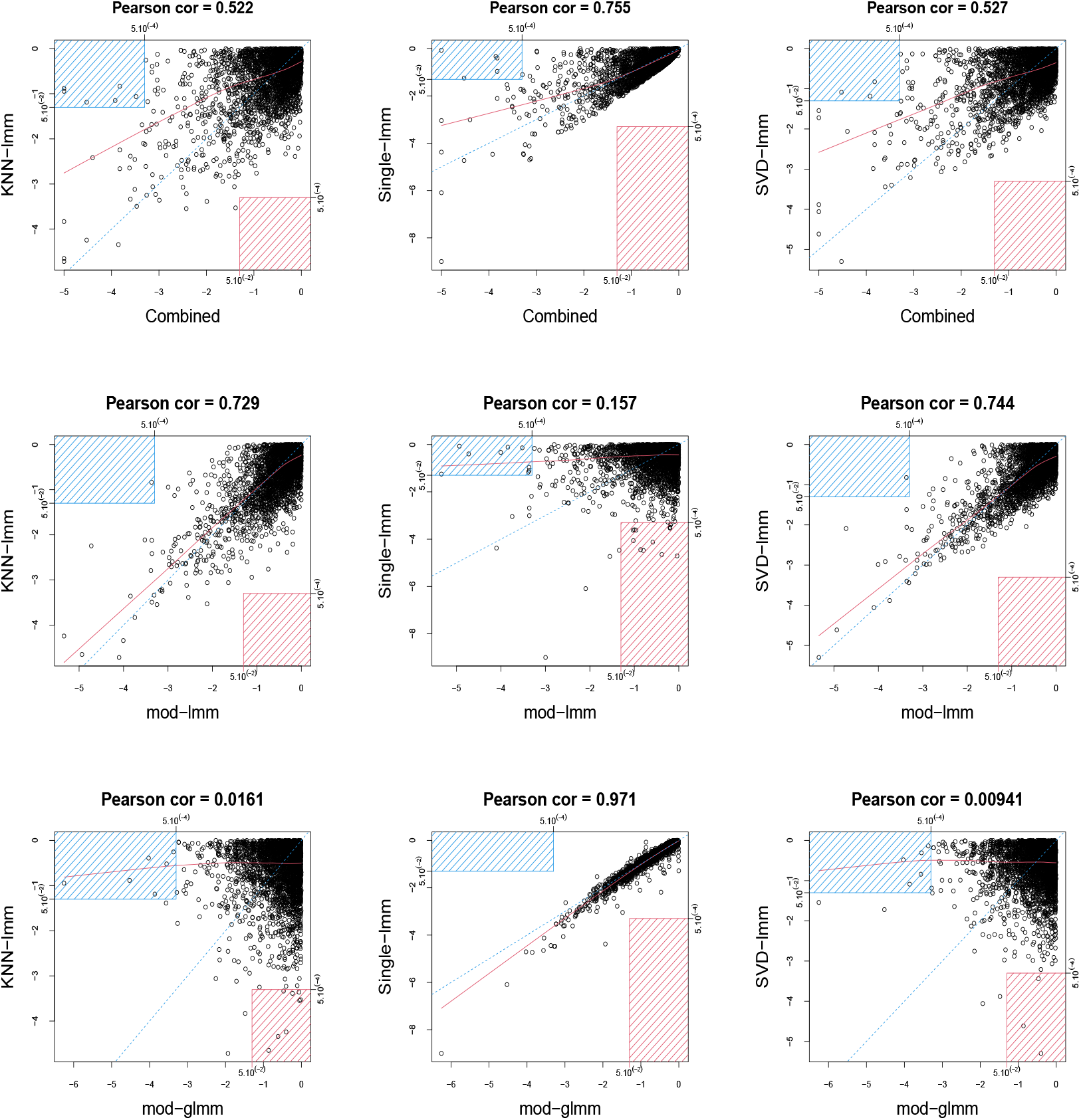
Scatterplots between log10-transformed *p*-values of pairs of FSMs for *Proteocardis-env*. Row 1: combined test and imputation-based FSMs. Row 2: Generalised mixed model (logistic) on missingness and imputation-based FSMs; proteins with less than 2 non-missing values are not displayed. Row 3: Linear mixed model on observed values and imputation-based FSMs. For each pair of testing procedure, the red rectangle corresponds to proteins with *p* > 5.10^-2^ with the first procedure and with *p* < 5.10^-4^ for the second procedure; conversely, the blue rectangle corresponds to proteins with *p* < 5.10^-4^ with the first procedure and with *p* > 5.10^-2^ for the second procedure.

**Figure S5.**
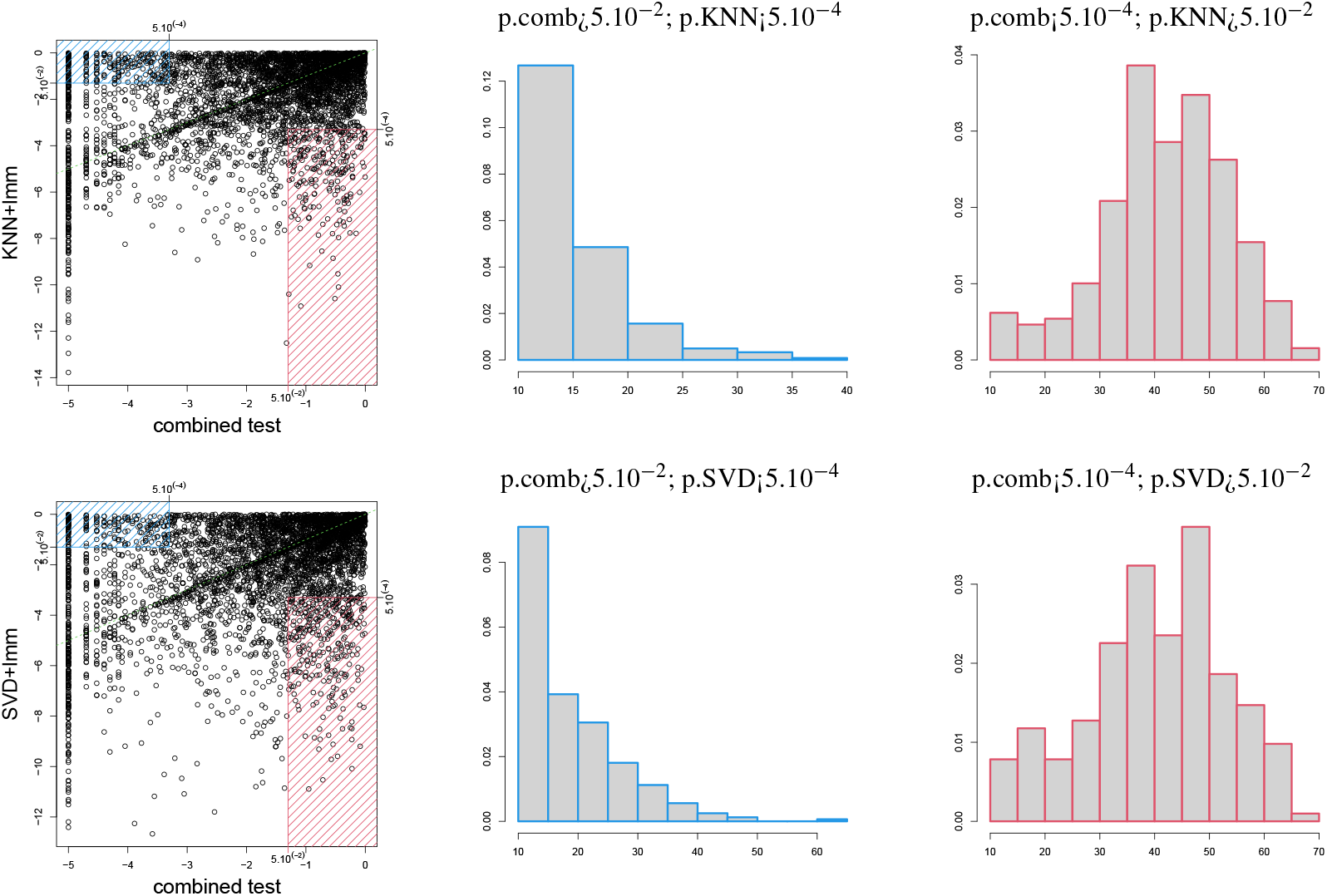
**Sparsity for proteins which are discordant** between the combined test and KNN-lmm (first row) or SVD-lmm (second row), on *Pigs*. Column 1: scatterplot of log10-transformed *p*-values of pairs of FSMs; the red rectangle corresponds to proteins with *p* > 5.10^-2^ with the first procedure and with *p* < 5.10^-4^ for the second procedure; conversely, the blue rectangle corresponds to proteins with *p* < 5.10^-4^ with the first procedure and with *p* > 5.10^-2^ for the second procedure. Column 2 (resp. 3): Histogram of the number of observed values by protein, for all proteins in the blue (resp. red) rectangle.

**Figure S6.**
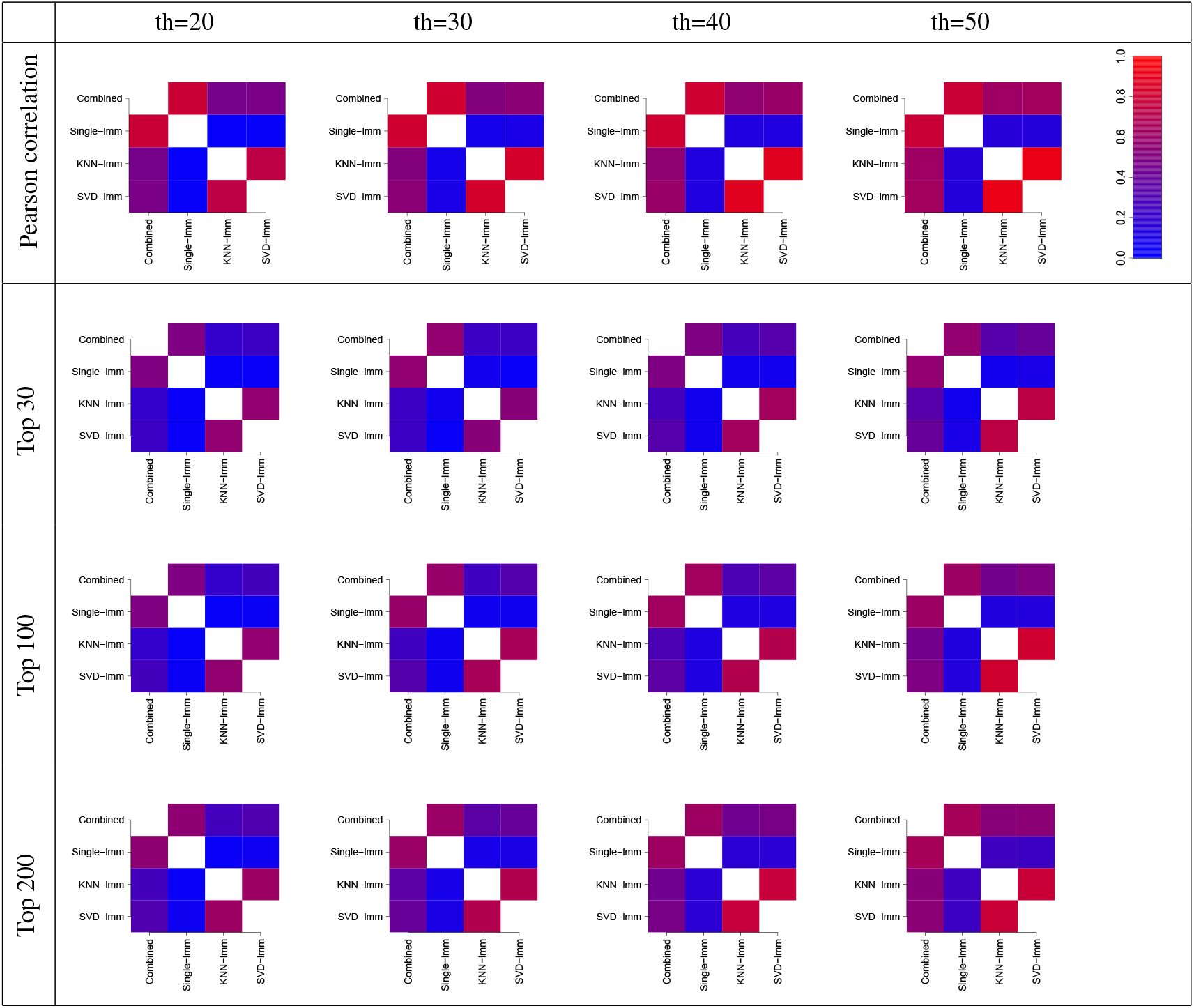
Pairwise agreement between *p*-values from the four FSMs, for filtering threshold of 20, 30, 40 and 50 for *Proteocardis-cyto*. Each row correspond to a criterion; row 1: Pearson correlation between log-transformed *p*-values; rows 2 to 4: proportion of common variables among the top *N* variables with *N* = 30, 100, 200. Each column correspond to a threshold value.

**Figure S7.**
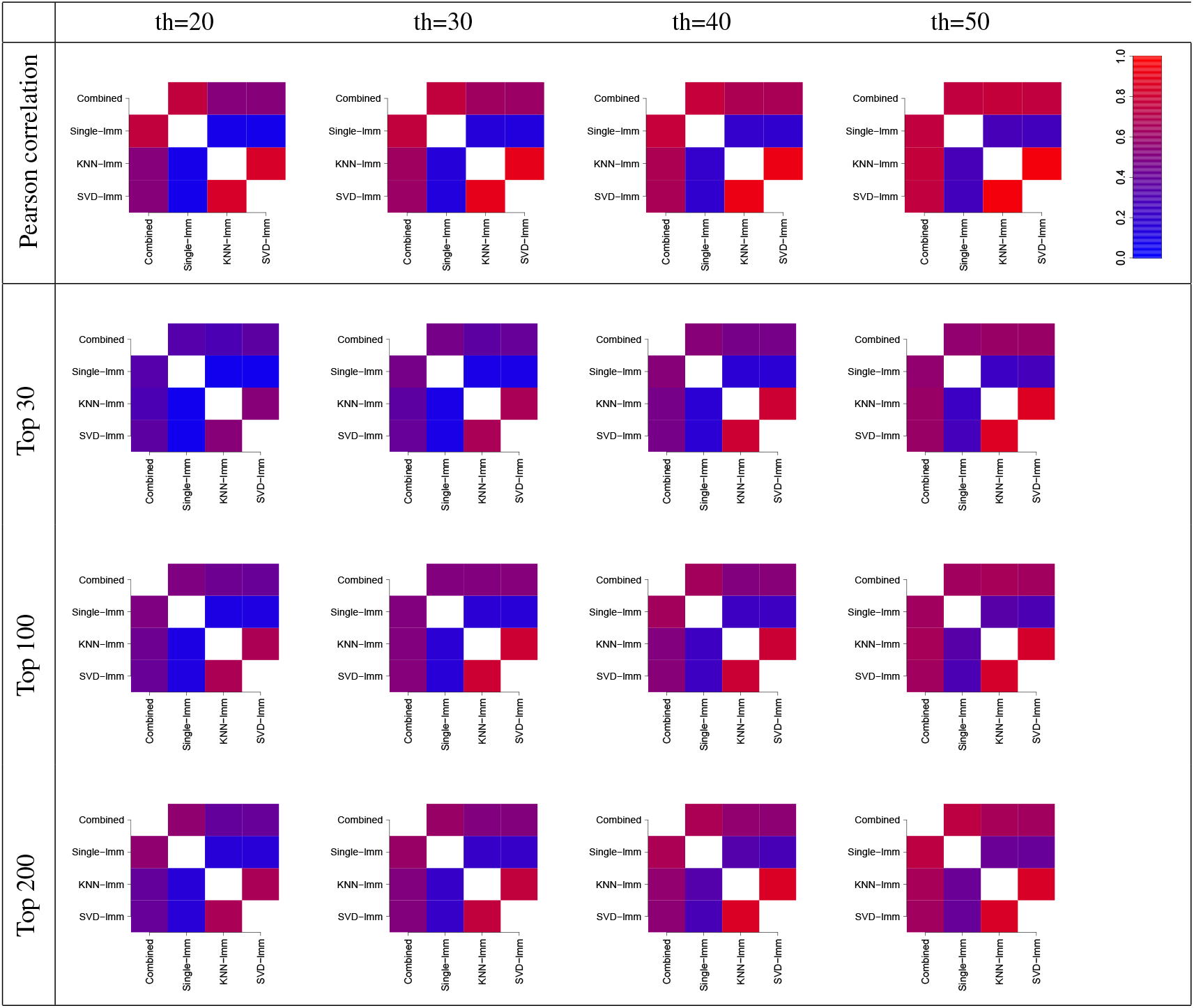
Pairwise agreement between *p*-values from the four FSMs, for filtering threshold of 20, 30, 40 and 50 for *Proteocardis-env*. Each row correspond to a criterion; row 1: Pearson correlation between log-transformed *p*-values; rows 2 to 4: proportion of common variables among the top *N* variables with *N* = 30, 100, 200. Each column correspond to a threshold value.

**Figure S8.**
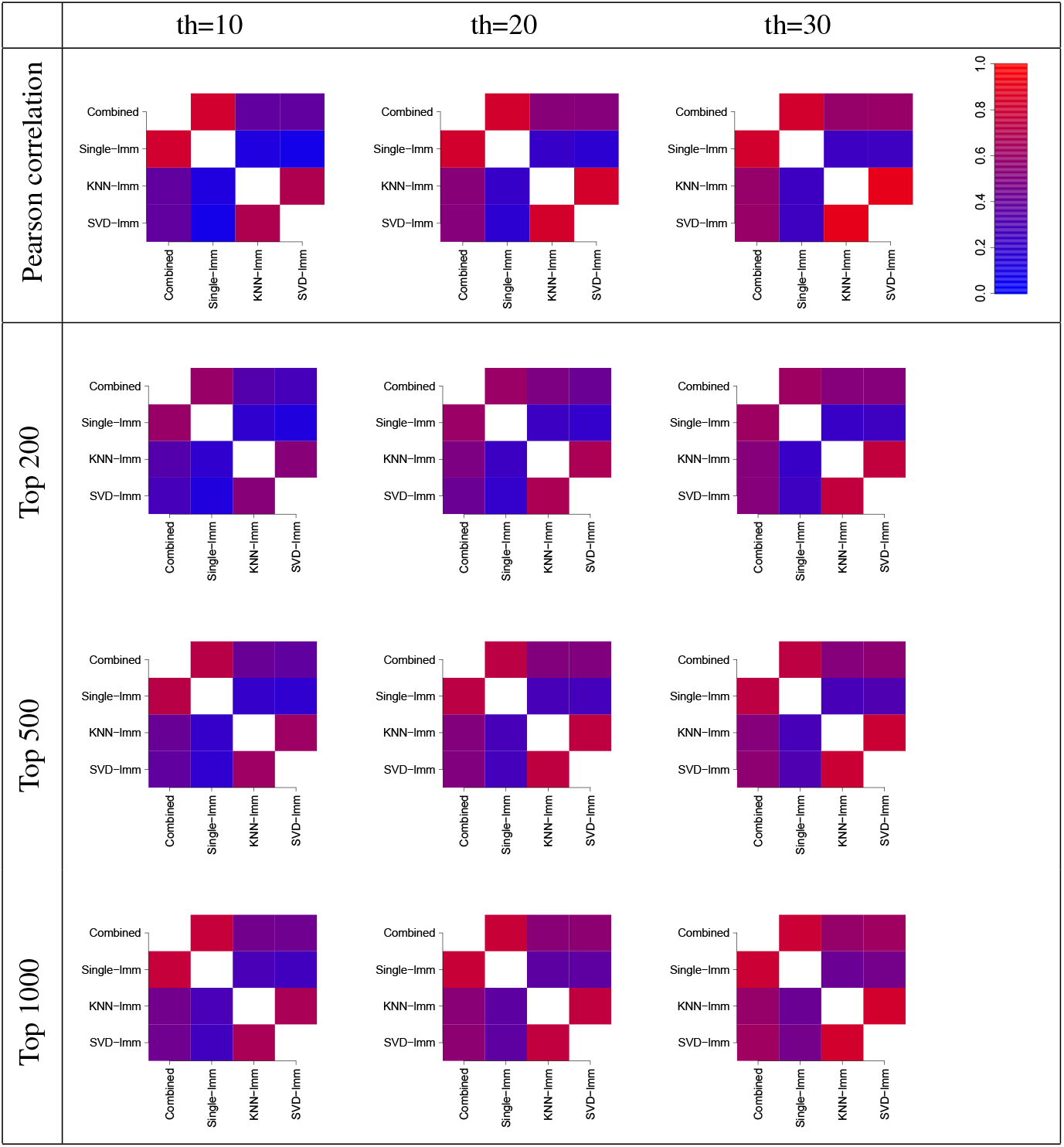
Pairwise agreement between *p*-values from the four FSMs, for filtering threshold of 20 and 30 for *Pigs*. Each row correspond to a criterion; row 1: Pearson correlation between log-transformed *p*-values; rows 2 to 4: proportion of common variables among the top *N* variables with *N* = 200, 500, 1000. Each column correspond to a threshold value.

**Figure S9.**
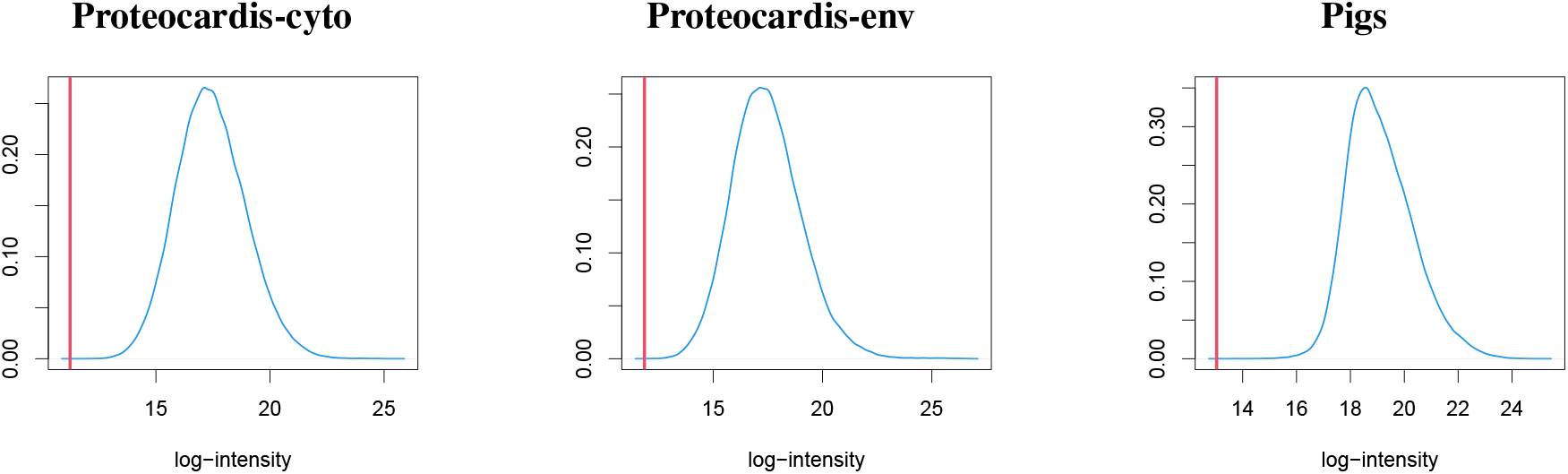
Single value imputation. Distribution of observed log-transformed intensities (blue) and imputed value (red) with single value imputation.

**Table S2.**
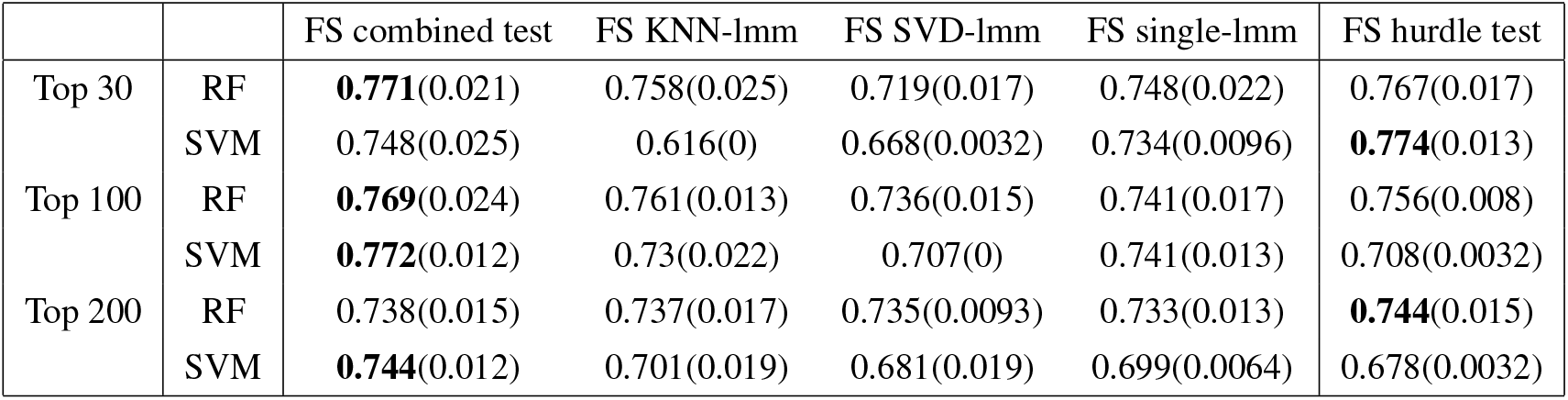
**Prediction accuracy** for two classification procedures on *Proteocardis-env*. The selection of the top *N* variables (*N* = 30, 100, 200) was followed by SVM or RF. Accuracy was computed in a 10-fold cross validation loop, repeated 10 times. Each cell provides the average accuracy (standard deviation of accuracy) computed over the 10 repetitions of the cross-validation. Bold numbers correspond to the highest accuracy among the four FSMs

**Figure S10.**
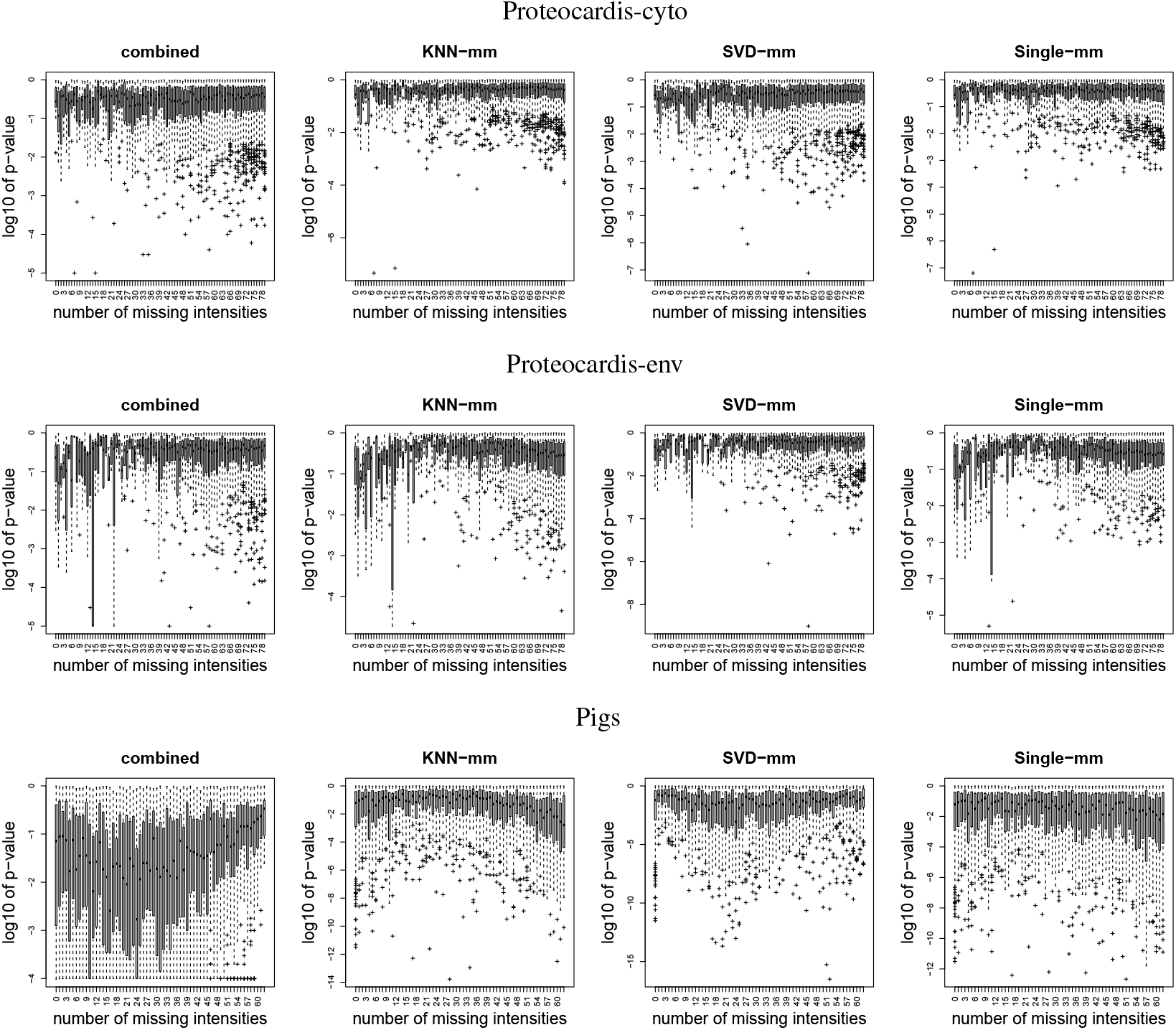
Log10-transformed *p*-values as a function of sparsity. The x-axis corresponds to the number of missing values among the 99 samples for *ProteoCardis* data sets, and among the 72 samples for *Pigs*.

**Figure S11.**
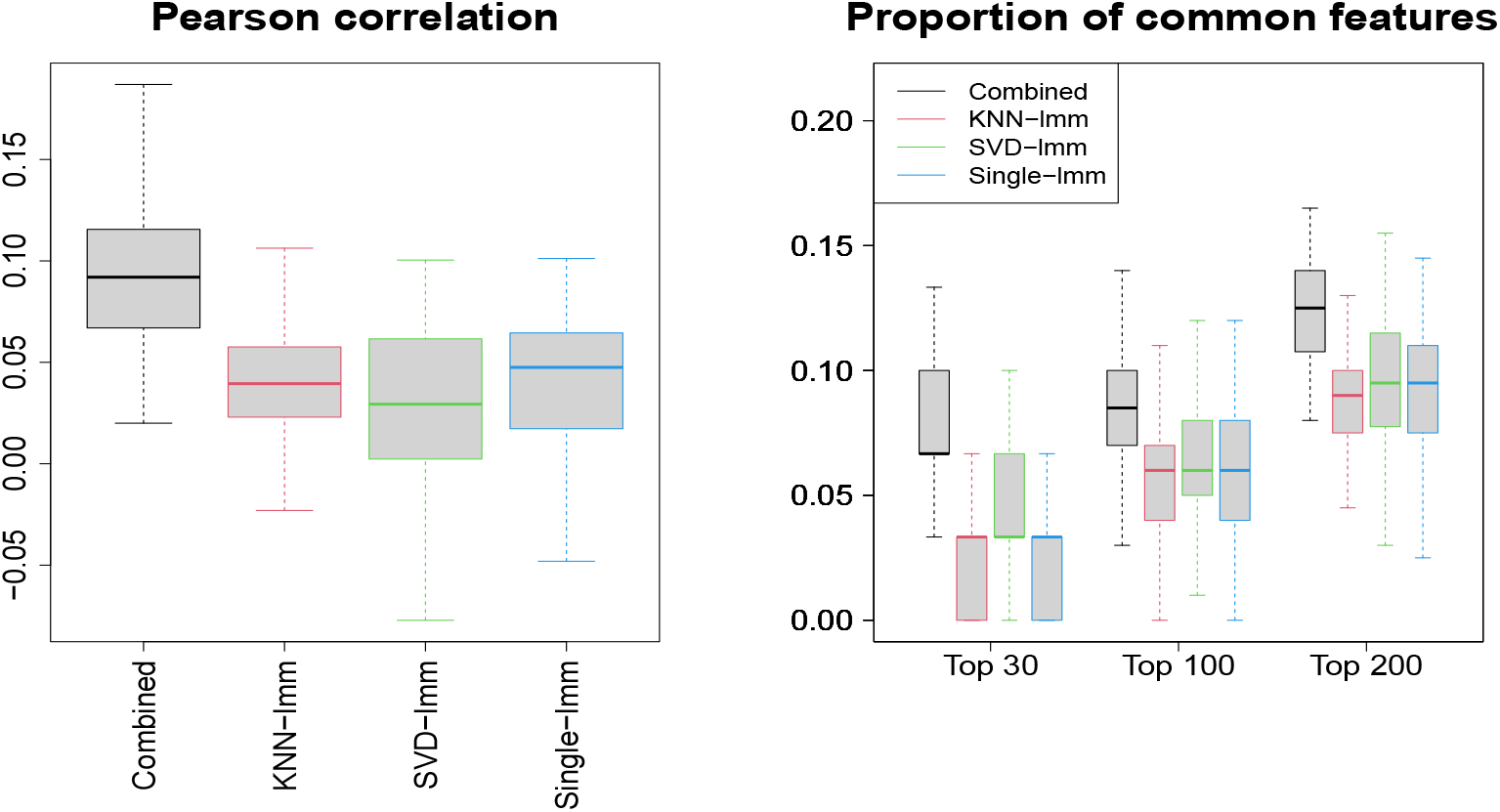
Replicability of variable selection on independent subsets. Pearson correlation between log-transformed *p*-values and proportion of common variables among the top *N* for 100 splitting of samples into two subsets. Dataset: *Proteocardis-env*.

**Figure S12.**
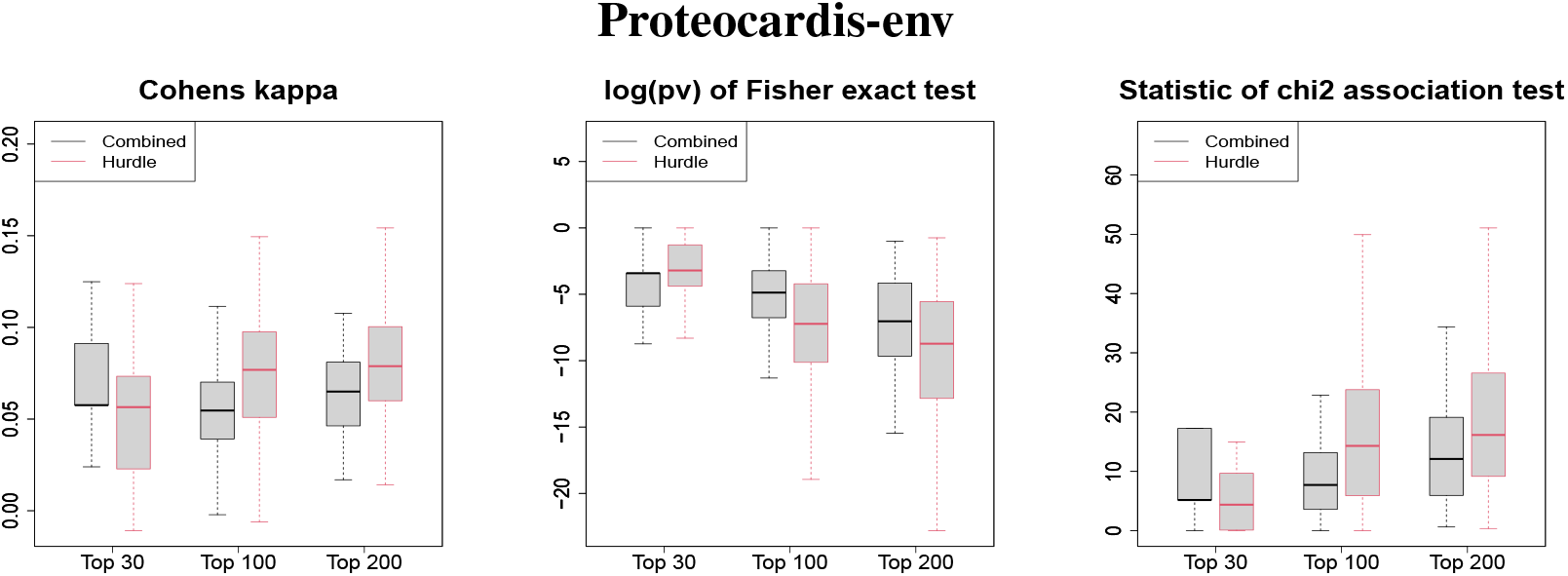
Replicability of variable selection on independent subsets for the hurdle test and the combined test. Boxplot of the Cohen’s kappa (left), the log-transformed *p*-value of Fisher test (center) and the statistic of the *χ*^2^ contingency table test (right), for selection of the top *N* features, performed on 100 splitting of the samples into two subsets. Black and red boxlots correspond to feature selection with the combined and the hurdle test respectively. Dataset: *ProteoCardis-env*

